# Protein interaction network analysis of mTOR signaling reveals modular organization

**DOI:** 10.1101/2023.08.04.552011

**Authors:** Devin T. Wehle, Carter S. Bass, Josef Sulc, Ghayda Mirzaa, Stephen E.P. Smith

## Abstract

The mammalian target of rapamycin (mTOR) is a serine-threonine kinase that acts as a central mediator of translation, and plays important roles in cell growth, synaptic plasticity, cancer, and a wide range of developmental disorders. The signaling cascade linking lipid kinases (PI3Ks), protein kinases (AKT) and translation initiation complexes (EIFs) to mTOR has been extensively modeled, but does not fully describe mTOR system behavior. Here, we use quantitative multiplex co-immunoprecipitation to monitor a protein interaction network (PIN) composed of 300+ binary interactions among mTOR-related proteins. Using a simple model system of serum deprived or fresh-media-fed mouse 3T3 fibroblasts, we observed extensive PIN remodeling involving 27+ individual protein interactions after one hour, despite phosphorylation changes observed after only five minutes. Using small molecule inhibitors of PI3K, AKT, mTOR, MEK and ERK, we define subsets of the PIN, termed ‘modules’, that respond differently to each inhibitor. Using primary fibroblasts from individuals with overgrowth disorders caused by pathogenic *PIK3CA* or *MTOR* variants, we find that hyperactivation of mTOR pathway components is reflected in a hyperactive PIN. Our data define a “modular” organization of the mTOR PIN in which coordinated groups of interactions respond to activation or inhibition of distinct nodes, and demonstrate that kinase inhibitors affect the modular network architecture in a complex manner, inconsistent with simple linear models of signal transduction.

## Introduction

Signal transduction networks have traditionally been modeled as linear cascades, where one protein acts upon the next in an orderly series of molecular events. Decades of research on mTOR signaling have demonstrated that signal-induced translocation of phosphoinositide 3-kinase (PI3K) phosphorylates membrane inositol PIP2 to PIP3, leading to AKT phosphorylation by the kinase PDK1 and the mTOR-containing protein complex mTORC2 (reviewed in Liu and Sabatini, 2020). Activated AKT phosphorylates TSC2, leading to TSC2 degradation and the release and disinhibition of the small GTPase Rheb. Rheb then activates mTORC1 by binding distally from the kinase site and causing a conformational change that accelerates catalysis and functionally activates mTOR kinase activity (Yang et al., 2017). Activated mTORC1 promotes initiation of cap-dependent translation and protein synthesis through the phosphorylation/activation of two related proteins, S6K1 and EIF4E binding protein (4EBP), which promotes the formation of the ribosomal complexes that initiate translation (Brunn et al., 1997; Burnett et al., 1998). In this manner, mTOR regulates the translation of hundreds of mRNA targets with a wide array of cellular functions (Masvidal et al., 2017), making mTOR a critical regulator of the life cycle of eukaryotic cells (Fig S1A).

While linear depictions of signaling networks can be conceptually useful by illustrating a hierarchy of kinase signaling, they do not accurately reflect the underlying molecular interactions of the protein interaction network (PIN); biology is not linear (Fig S1B,C). Rather, the protein components of signaling networks interact to form a complex network topology that includes feed-back or feed-forward loops, or cross-talk between seemingly unrelated pathways (Mendoza et al., 2011). For example, PI3K-dependent membrane phosphorylation potentially recruits 247 human proteins that contain a plextrin homology (PH) domain (Rusten and Stenmark, 2006) (although only perhaps 10% of PH-domain containing proteins bind phosphoinositide (Lemmon, 2007)). AKT affects the phosphorylation of 100s of targets (Manning and Cantley, 2007; Wiechmann et al., 2021), including GSK3B and mSin1, an obligate component of mTORC2 (Humphrey et al., 2013). Moreover, proteins that are primarily thought of as acting at the “top” of the signaling cascade physically interact with proteins at the ‘bottom’ of the pathway, e.g. PDK1 phosphorylates p70S6 kinase directly (Pullen et al., 1998). Thus, conceptualizing mTOR signaling as a sequence of phosphorylation events may give an incomplete view of how the signal transduction network functions.

An alternative conceptual framework for modeling signal transduction relies on protein interaction networks, whose network topography is acutely post-translationally modified during signaling events. In response to environmental stimuli, coordinated groups of protein-protein interactions change their co-associations in unison, modifying the structure and function of the cellular interactome (Heavner et al., 2021; Lautz et al., 2021, 2018; Lundby et al., 2019; Smith et al., 2016). The dynamic composition of these protein complexes (Pawson and Nash, 2003), and the magnitude of their activation (Neier et al., 2019), is thought to instruct specific cellular responses. Different stimuli may engage different groups of coordinated interactions, termed “modules” (Coba et al., 2009; Lautz et al., 2018; Pawson and Nash, 2003), or engage identical modules of interactions with different intensities or kinetics (Neier et al., 2019), allowing the cell to disambiguate multiple environmental cues by activating signal transduction networks in subtly different ways. Here, for the first time, we characterize a dynamic protein interaction network composed of mTOR-related signaling proteins by activating or inhibiting different nodes of the mTOR pathway and measuring the PIN response using quantitative multiplex co-immunoprecipitation (QMI). Our data reveal a complex network structure with extensive interactions among ‘upstream’ and ‘downstream’ components and demonstrate an unexpected modular organization in which PI3K and mTOR inhibition act on distinct groups of binary interactions spanning all levels of the linear mTOR hierarchy.

## RESULTS

### mTOR target selection and assay development

QMI is an antibody-based approach that measures dynamic changes in protein interaction networks by simultaneous immunoprecipitation of multiple protein targets onto flow cytometry beads, followed by simultaneous detection using a set of fluorescently labeled “probe” antibodies that bind to different epitopes (Fig 1A). The median fluorescent intensity (MFI) of each IP_Probe pair is used as a proxy for the magnitude of the interaction between the IP antibody target and the probe antibody target. Experiments are run on carefully matched pairs of samples that are identical except for a recent experimental manipulation; any differences between the IP-probe data matrices are assumed to be due to acute changes in protein complexes caused by the manipulation. Extensive internal controls account for background fluorescence and differences in bead distributions between replicates, and correct for batch-to-batch variability (Brown et al., 2019; Smith et al., 2016). Critically, because multiple binary interactions are measured simultaneously, we can use network approaches such as weighted correlation network analysis (Langfelder and Horvath, 2008) to model the behavior of multiple interactions changing in unison, which may reduce biological noise associated with single-protein measurements and reveal modules of interactions that encode features of the experimental manipulation (Brown et al., 2019; Lautz et al., 2018; Smith et al., 2016).

**Figure 1:**
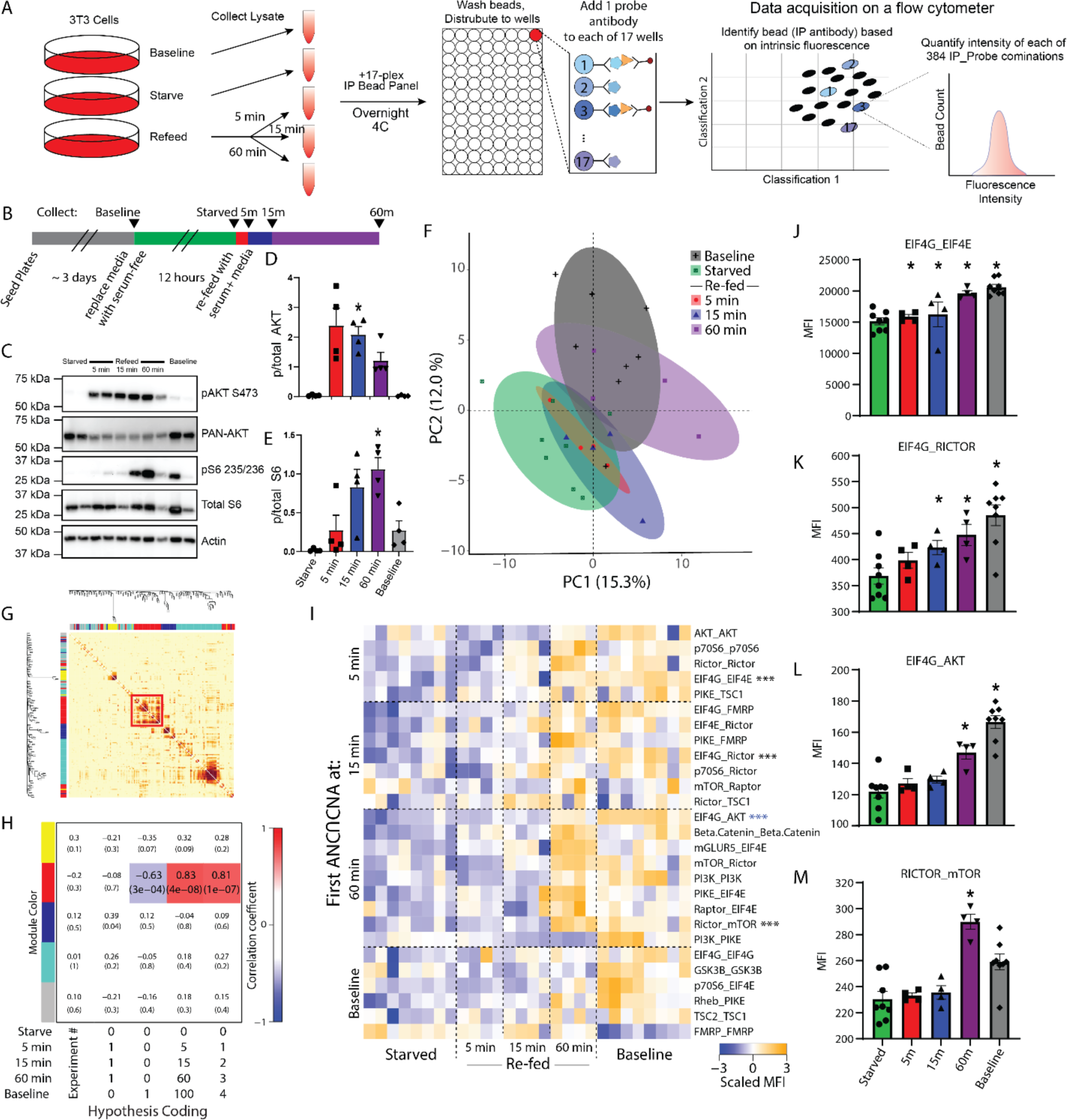
Kinetics of mTOR protein network dynamics. A) Quantitative multiplex co-immunoprecipitation (QMI) procedure. See methods for details. B) Experimental design. C) Representative western blots showing phospho- and total AKT and S6, with actin for a loading control. C-E) Quantification of blots shown in B. * indicates significantly different from starved by ANOVA followed by Bonferroni-correction post-hoc testing, p<0.05. N = 4. F) Principal component graph of QMI data, N=4-8. G) Topological overlap matrix (TOM) clusters interactions into modules as indicated by the colored rectangles at top and left. Each pixel indicates the level of overlap between the interactions in the corresponding row and column, red color indicates high overlap. The significant “red” module is bounded by a red box. H) Module-trait table showing the correlation coefficient (top number) and p-value (bottom number) between the eigenvector of each color-coded module (colored rectangles at left) and binary-coded trait labels (“hypotheses”) shown in the table below. I) Heatmap of the median scaled values of all significantly altered interactions. Each box represents a single interaction measurement from a single biological replicate; columns correspond to a biological replicate while rows correspond to an interaction (IP_Probe), ordered by the timepoint that the interaction first reached significance. N = 4-8. Statistical significance calculated by ANC and CNA statistics as detailed in methods, only interactions significant by both tests are listed. *** indicates an interaction that is being represented by subsequent bar graphs. J-L) Bar graphs showing the median fluorescent intensity (MFI) of representative interactions, indicated with asterisks in H. * indicates significantly different from starved by ANC, a paired statistical test designed for QMI data.

Targets for the mTOR antibody panel were selected based on four criteria: known relevance to mTOR signaling (Fig S1A), known co-associations in at least one protein complex, as listed in BioGrid, IntACT or specific literature searches (Fig S1B); known association with autism spectrum disorder using the SFARI Gene database (queried in 2018); and the existence of suitable, commercially available antibodies. We screened 3-5 antibodies for each target to identify two different antibodies that could simultaneously bind in the native state, and validated antibody specificity in immunoprecipitation-flow cytometry using knockout or overexpression strategies (detailed in Fig S2, S3 and Table S1) to identify antibody pairs that specifically recognized 17 members of the mTOR PIN in mouse and human. Since lysis buffer detergent has a strong effect on which interactions are detected (Lautz et al., 2019), we optimized detergent conditions and determined 1% Digitonin identified the largest number of interactions (Fig S4).

### mTOR PIN kinetics following starvation/refeeding

To establish a basic framework for QMI measurement of mTOR PIN dynamics, we serum-starved NIH 3T3 cells for 12 hours to dampen constitutively active mTOR signaling, then stimulated by re-feeding with fresh media for 5, 15 or 60 minutes before lysis; controls included cells that were never starved (baseline) or starved cells that were never re-fed (Fig 1A-B). Western blots showed the expected pattern of AKT phosphorylation peaking at 5 minutes and gradually diminishing over the subsequent timepoints, while phospho-S6 increased at 15 min and remained elevated at 60 min (Fig 1C-E). QMI was performed on cell lysates, resulting in a data matrix consisting of ∼50-150 individual flow cytometry bead reads for each of 384 binary interactions measured, in duplicate, for each of 28 experimental total measurements, for over 2.1 x 10^6^ total bead reads. The bead distributions were collapsed into a single median fluorescent intensity value (MFI) for each IP_Probe, and principal component analysis (PCA) was used to visualize the overall data structure. Starved cells clustered separately from baseline (Fig 1F), but, contrary to western blot data, after 5 or 15 minutes of re-feeding samples still overlapped with the starved condition, only resembling the baseline condition after 60 min. Correlation network analysis (Langfelder et al., 2008) of the same MFI dataset identified a single module of interactions whose behavior correlated with other interactions in the module across the 28 experimental replicates, as visualized by a topological overlap matrix (TOM) plot (Fig 1G). The eigenvector defining the overall behavior of this module, arbitrarily color-coded “red” by the analysis program, showed that its behavior was best correlated with a binary-coded hypothesis of “time since refeeding” (correlation coefficient = 0.83, p = 4 x 10^-8^) (Fig 1H), indicating a slow increase in the median scaled value of the module. No other module correlated with any experimental variables, causing us to focus on this “red” module.

To ensure reporting of only robust, high-confidence interactions, we performed a second adaptive, non-parametric statistical test corrected for multiple comparisons (ANC, see methods). ANC does not collapse bead distributions, but instead incorporates features of the distributions and the consistency of replicates to establish interactions that change in >70% of pair-wise comparisons at a multiple-comparison-corrected P<0.05 for each interaction across the entire experiment (Brown et al., 2019; Smith et al., 2016). ANC-significant interactions that were also significantly correlated with the “red” module were considered high-confidence interactions identified by both independent tests. Twenty-seven such interactions were visualized as a row-normalized heatmap, with rows sorted by the time at which the Interaction first became significant and columns arranged by re-feeding time (Fig 1I). At five minutes, two interactions slightly increased in abundance-EIF4G_EIF4E (Fig 1J) and PIKE_TSC1 (log_2_ Fold Change (log_2_FC) = 0.24 and 0.19, respectively) and were significant by ANC and CNA analysis. Additionally, the apparent abundance of three targets was reduced: AKT_AKT, p70S6_ p70S6 and RICTOR_ RICTOR. Changes in apparent abundance may reflect rapid proteasomal degradation, or a change in binding partners or a post-translational modification that occludes IP or probe antibody binding. Regardless, it was striking that, despite AKT phosphorylation clearly peaking at 5 minutes (Fig 1D), the overall network at 5 minutes was essentially unchanged. By 15 minutes, the two aforementioned interactions continued to increase (log_2_FC = 0.11 and 0.44, respectively), and 7 other interactions including EIF4G_RICTOR (Fig 1K), EIF4E_RICTOR and PIKE_FMRP began to increase (log_2_FC = 0.28, 0.25 and 0.31, respectively). By 60 min, these interactions further increased, and 9 additional interactions reached significance, including EIF4G_AKT (Fig 1L), RICTOR_mTOR (Fig 1M) and RAPTOR_EIF4E (log_2_FC = 0.39, 0.37 and 0.38, respectively). These data represent a core network of dynamic protein interactions that acutely change following mTOR activation.

### Inhibition of the mTOR pathway

To explore the relationship between a traditional linear model of mTOR signal transduction and our QMI approach, we starved cells and re-fed for 60 minutes while including one of three inhibitors of the mTOR pathway that act at different levels of the linear signaling model (Fig 2A). BKM120, which inhibits all four catalytic isoforms of class I PI3K in an ATP-competitive manner but does not inhibit mTOR (Maira et al., 2012), prevented the phosphorylation of both AKT and S6 by western blot, as expected (Fig 2B-C). RAD001 (Everolimus), which inhibits mTORC1 by forming a complex with FKBP-12 and partially occluding substrate entry into the mTORC1 active site (Yang et al., 2013), prevented S6 phosphorylation but not AKT phosphorylation, as expected. AZD5363, an ATP-competitive pan-AKT inhibitor (which also inhibits PKA and P70S6K) (Davies et al., 2012), caused an apparent increase in AKT phosphorylation and inhibited S6.

**Figure 2:**
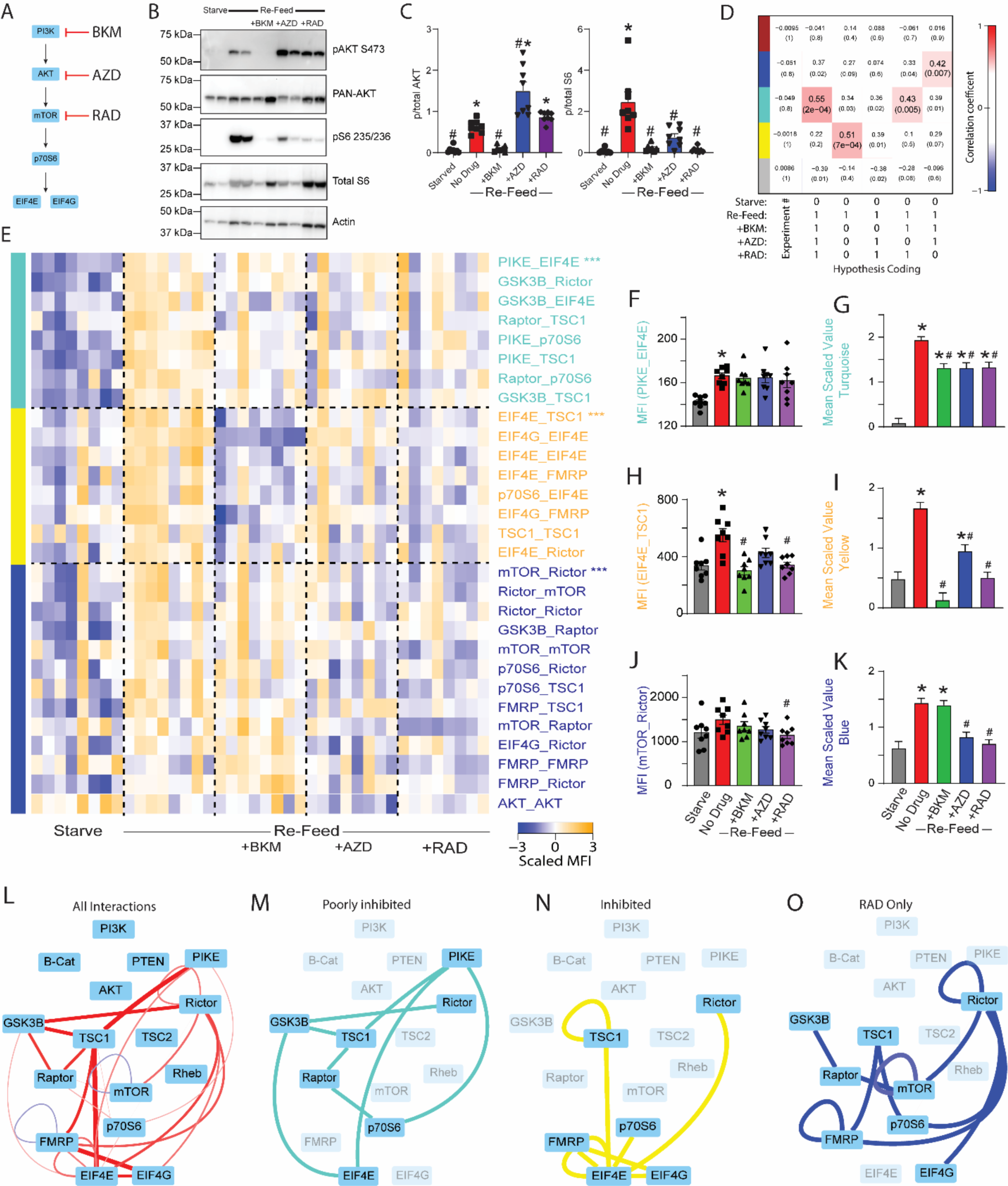
Inhibitors of PI3K/AKT/mTOR reveal modular pathway organization. A) Small molecule inhibitors of PI3K/AKT/mTOR were applied during a starve-refeed experiment. B) Western blots of phospho- and total AKT and S6, and actin for loading control, in 3T3 fibroblasts that were starved and re-fed in the presence of mTOR drugs. C) Quantification of blots in B. * indicates significantly different from starved, # indicates significantly different from re-fed by ANOVA followed by Bonferroni-corrected post-hoc testing, p<0.05. N=4. D) Module-trait table showing the correlation (top number) and the p-value (bottom number) between the eigenvector of each color-coded module (colored rectangles at left) and binary-coded trait labels (“hypotheses”) shown in the table below. E) Heatmap of the median scaled values of all significant interactions. Each box represents a single interaction measurement from a single biological replicate; columns correspond to a biological replicate while rows correspond to an interaction. Interactions are listed in descending order by module membership, color-coded by rectangles at left and by text color. N = 4-8. F) Median fluorescent intensity (MFI) of the interaction between PIKE_EIF4E, the interaction most correlated to turquoise module behavior. G) Mean scaled value of all interactions in the turquoise module, which corresponds to overall module behavior. H-K) Similar to F-G, but for yellow and blue modules. For F-K, * indicates significantly different from starved, # significantly different from re-fed, by ANOVA followed by Bonferroni-corrected post-hoc testing, p<0.05. L) Dynamic interactions that showed significant changes with re-feeding are represented by lines connecting protein nodes. Red indicates that the interaction was increased following re-feeding, blue indicates decreased; the thickness of the line indicates relative magnitude of the change. M) Interactions in the turquoise module that was partially inhibited by all drugs. N) Interactions in the yellow module that was inhibited by all drugs, most strongly by BKM. O) Interactions in the blue module that was inhibited by RAD and AZD, but not by BKM.

Following QMI, we independently re-calculated modules based on the new dataset using correlation network analysis. The inclusion of inhibitors in the experiment caused the “red” module identified in Figure 1 to fracture into three modules that correlated with different binary-coded hypotheses (Fig 2D). First, a module color-coded “turquoise” most strongly correlated with the hypothesis of activation across all re-fed conditions, regardless of drug (correlation coefficient = 0.55; p = 0.0002). Next, a module coded “yellow” most significantly correlated with activation during refeeding and inhibition by *all* drugs (correlation coefficient = 0.51; p = 0.0007). Finally, a module coded “blue” most strongly correlated with activation by refeeding and inhibition by RAD only (correlation coefficient = 0.42; p = 0.007). Again, interactions correlated with each module were merged with interactions identified by the ANC statistical test, and twenty-nine significant interactions were visualized as a row-normalized heatmap, with rows ordered by module membership and columns by treatment. The “turquoise” module was activated by refeeding and partially (although not completely) inhibited by all three drugs (Fig 2E, top), as demonstrated by the behavior of the protein interaction most highly correlated to module behavior, PIKE_EIF4E (Fig 2F), and by the mean scaled value of all interactions in the module (Fig 2G). The yellow module, exemplified by EIF4E_TSC1 (Fig 2H), was increased by re-feeding, and inhibited by all drugs, most strongly BKM (Fig 2I). Finally, a blue module, exemplified by mTOR_Rictor (Fig 2J) was significantly inhibited by RAD and AZD, but surprisingly not by BKM (Fig 2K).

To visualize the entire network, we represented all significant interactions in a node-edge diagram, where the thickness/color of the edges indicates the magnitude/direction of the change in each binary measurement (Fig 2L), as well as the subset of interactions that comprise each module (Fig 2M-O). The poorly inhibited “turquoise” module included proteins involved at all levels of mTOR signaling, including GSK3B, PIKE, PICTOR, p70S6 and EIF4E. Partial activation of this module in the presence of all inhibitor drugs may reflect crosstalk with other signaling pathways that respond to refeeding, including GSK3B or ERK activity, which were not manipulated in this experiment. The “blue” (RAD-responsive) module contained interactions including mTOR, RAPTOR, and RICTOR that one might have expected to be strongly inhibited by RAD, including mTOR_RAPTOR. This module was inhibited by AZD and RAD, but surprisingly, not by BKM, suggesting other parallel inputs may affect this module and can compensate for the loss of PI3K signaling. The “yellow” (inhibited) module contained the most downstream interactions, including EIF4E, EIF4G, p70S6 and FMRP. These interactions were significantly inhibited by all inhibitors, but the magnitude of inhibition was largest for BKM>RAD>AZD. In fact, AZD still permitted significant module activation. It is somewhat surprising that ‘upstream’ PI3K inhibition was so effective at targeting these ‘downstream’ interactions, despite BKM’s inability to inhibit the blue RAD-dependent module. Overall, these data demonstrate an unexpectedly complex modular organization of the mTOR PIN.

The reduction in both mTOR_Rictor and mTOR_Raptor by RAD was unexpected, since short-term treatment with RAD is thought to affect only mTORC1, and phospho-western blots showed the expected lack of effect of RAD on AKT phosphorylation (Fig 2C). We therefore performed co-immunoprecipitation-western blot assays on 3T3 fibroblasts treated with RAD, and confirmed a reduction of both IP:mTOR western blot:Raptor and IP:mTOR western blot:Rictor following RAD treatment (Figure S5). These data are consistent with previous reports showing disruption of both mTORC1 and mTORC2 protein complexes, although not necessarily AKT phosphorylation, by Rapamycin (Rosner and Hengstschläger, 2008; Sarbassov et al., 2006).

### The contribution of the ERK Pathway

The failure of BKM to prevent changes in the RAD-responsive “blue” module suggested non-canonical routes of activation, i.e., crosstalk from other signal transduction pathways. The ERK/MAPK pathway can affect mTOR signaling via inhibition of the TSC complex, as well as multi-level feedback inhibition (Mendoza et al., 2011). To observe the contribution of Erk signaling to stimulus-dependent changes in the mTOR PIN, we performed a starve-refeed experiment in the presence of three different ERK pathway inhibitors (Fig 3A) and confirmed their efficacy by western blot (Fig 3 B-E). FR180204, a selective ERK1/2 inhibitor that binds to the ATP binding pocket of ERK (Ohori et al., 2007), increased ERK phosphorylation and eliminated S6 phosphorylation while having no effect on AKT phosphorylation. Both U0126, a selective inhibitor of MEK1 and MEK2 (Favata et al., 1998), and PD98059, which binds to the inactive form of MEK and prevents its phosphorylation by RAF-1 and downstream activity (Dudley et al., 1995), inhibited ERK phosphorylation, increased AKT phosphorylation, and partially inhibited p-S6 (although U0126 was less effective than PD98059). RAD was included to delineate mTOR-dependent interactions, and inhibited S6 phosphorylation as expected while having no effect on ERK.

**Figure 3:**
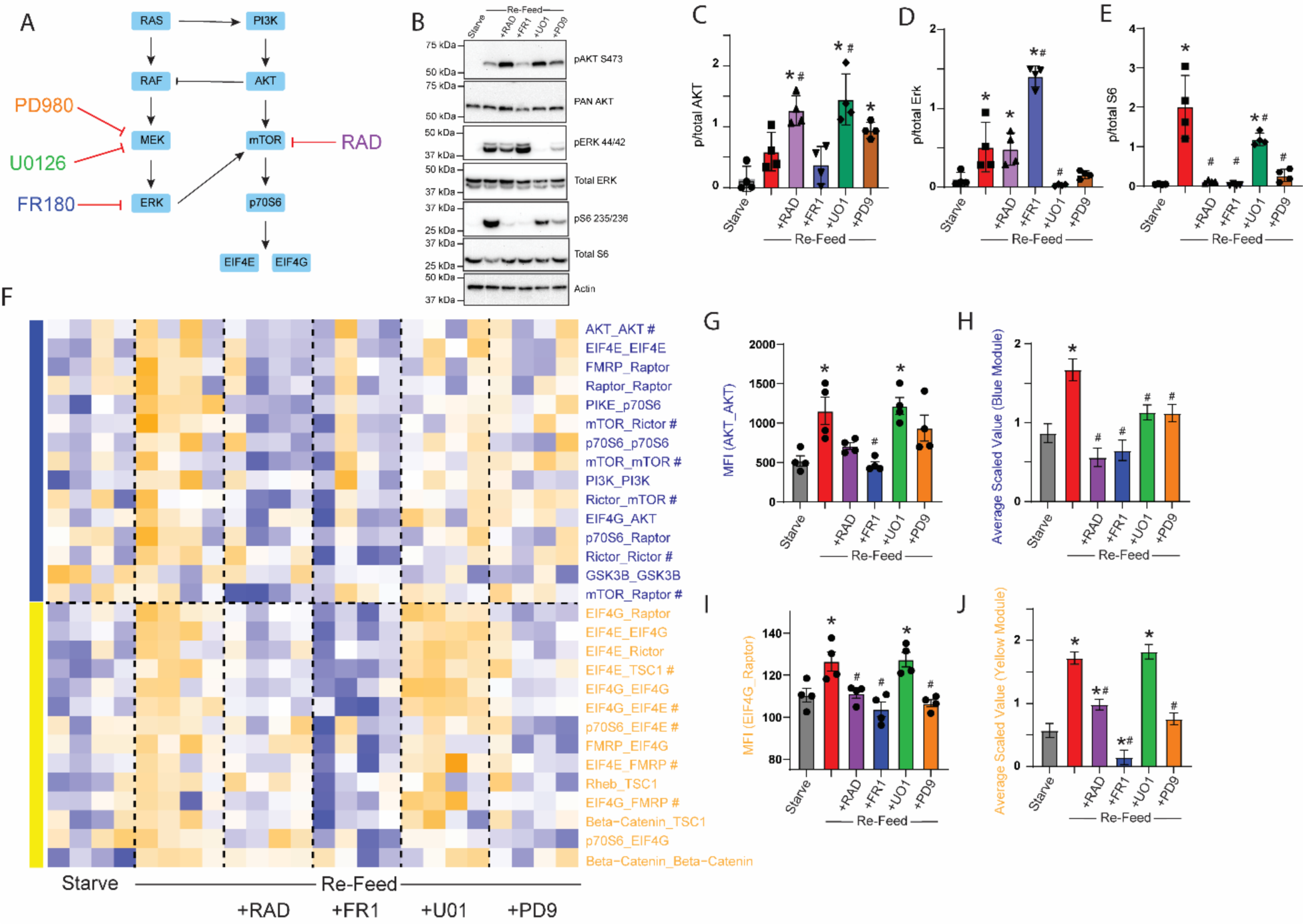
MEK and ERK inhibitors differentially affect mTOR modules. A) Small molecule inhibitors of MEK, ERK or mTOR were applied during a starve-refeed experiment. B) Western blots of phospho- and total AKT, ERK and S6, and actin for loading control, in 3T3 fibroblasts that were starved and re-fed in the presence of drugs. C-E) Quantification of blots in B. * indicates significantly different from starved, # indicates significantly different from re-fed by ANOVA followed by Bonferroni-corrected post-hoc testing, p<0.05. N=4. F) Heatmap of the median scaled values of all significant interactions changed with re-feeding. Each box represents a single interaction measurement from a single experiment; columns correspond to an experiment while rows correspond to an interaction. N = 4. Interactions are ordered by module membership, represented by colored bars on the left, and coloring of Interaction. G) Median fluorescent intensity (MFI) of AKT_AKT, the interaction most correlated with the blue module. H) Mean scaled value of all interactions in the blue (RAD-responsive) module, which corresponds to overall module behavior. I-J) Similar to G-H, but for the yellow module. For G-J, * indicates significantly different from starved, # significantly different from re-fed, by ANOVA followed by Bonferroni-corrected post-hoc testing, p<0.05.

Following QMI, we again independently re-calculated CNA modules and identified two that correlated with experimental variables (Fig 3F). A “blue” RAD-responsive module (which was calculated independently from the blue module in Figure 2 but contained similar interactions) was increased by re-feeding and strongly inhibited by both RAD and FR180 (Fig 3G). The MEK inhibitors U0126 and PD98059 had an intermediate effect, as demonstrated by both the interaction most strongly correlated to the blue module, AKT_AKT (Fig 3G) and by the average scaled value of the module (Fig 3H). MEK inhibition significantly inhibited interactions compared to the re-fed/no drug condition, but not as strongly compared with ERK or mTOR inhibition (although the difference between the starve/no drug and re-fed+drug conditions were not significant for any drug, Fig 3H). Of note, six interactions from the “blue” module that was independently identified in Figure 2, including mTOR_Raptor, were also included in this module. Overall, these data show strong inhibition of the blue module by both RAD and ERK inhibitors.

A second, yellow module (again derived independently from, but containing similar interaction to, the yellow module in Figure 2), contained a set of interactions that were inhibited in unexpectedly different ways by our drug panel. Exemplified by EIF4G_Raptor (Fig 3I), this module showed strong inhibition by FR180, which reduced activity significantly lower than the starved condition (Fig 3J). RAD and PD98059 significantly inhibited the activation of the yellow module compared to re-feeding alone, although RAD was still significantly increased over the starved state. Surprisingly, MEK inhibition with U0126 had no effect on yellow module activation. The yellow module contained many interactions related to the EIF4 translation initiation complex and included 6 (out of 8 total) of the BKM-sensitive yellow module interactions derived independently in Figure 2. Thus, direct ERK inhibition was similar to PI3K inhibition, while MEK inhibition produced strikingly different results: U0126 was ineffective, while PD98059 was as effective as RAD.

### Next Generation mTOR Inhibitors

Multiple generations of mTOR inhibitors have been created since the discovery of rapamycin. TORIN1 (TRN) and Sapanisertib (SAP) are both second-generation ATP competitive small molecule inhibitors that target mTOR in both mTORC1 and mTORC2 (Gökmen-Polar et al., 2012; Liu et al., 2010). Rapalink-1 (RLK) is a third generation mTOR inhibitor formed by physically linking rapamycin and SAP with an inert chemical linker that simultaneously forms an inhibitory complex with FKBP-12 and blocks the mTOR ATP binding site (Rodrik-Outmezguine et al., 2016). We repeated the starve/refeed paradigm in the presence of each inhibitor to directly compare their effects on the mTOR PIN.

We first determined equivalent doses of the four drugs, which we defined as the lowest concentration that inhibited phosphorylation of AKT (where applicable), RPS6, p70S6K, and 4EBP1 in a titration experiment using increasing concentrations of drug, starting at each drug’s IC-50 (Figure Fig S6A-D). Phospho-westerns showed that the selected doses of TRN and SAP inhibited AKT phosphorylation, indicating mTORC2 inhibition, while RAD and RLK did not (Figure 4A-B). As expected, both RPS6 and p70S6K1 were inhibited by all drugs (Figure 4C-D). However, only TRN and SAP affected 4EBP1 phosphorylation, consistent with previous studies (Liu et al., 2010; Rodrik-Outmezguine et al., 2016) (Figure 4E). QMI identified two modules that correlated with experimental variables: a yellow module that was most significantly correlated with activation during re-feeding and inhibition by all drugs (correlation coefficient 0.56; p = 0.0004) and a blue module that most significantly correlated with activation during re-feeding but inhibition by SAP (correlation coefficient -0.43; p = 0.003) or RLK treatment (correlation coefficient 0.41; p = 0.04) (Figure 4F). The thirty-two significant interactions identified by ANC and CNA were visualized as a row normalized heatmap (Figure 4G). The yellow module was composed of primarily EIF4E/EIF4G/p70S6 interactions, indicating changes in translation-related protein complexes downstream of mTOR. These interactions were increased by re-feeding, inhibited by RAD, and strongly inhibited by next generation mTOR inhibitors such that the averaged scaled value of the module was lower than in the starved condition (Figure 4H,I). These data demonstrate that next generation mTOR inhibitors are significantly more efficient at mTOR inhibition compared to either RAD or overnight serum starvation.

**Figure 4:**
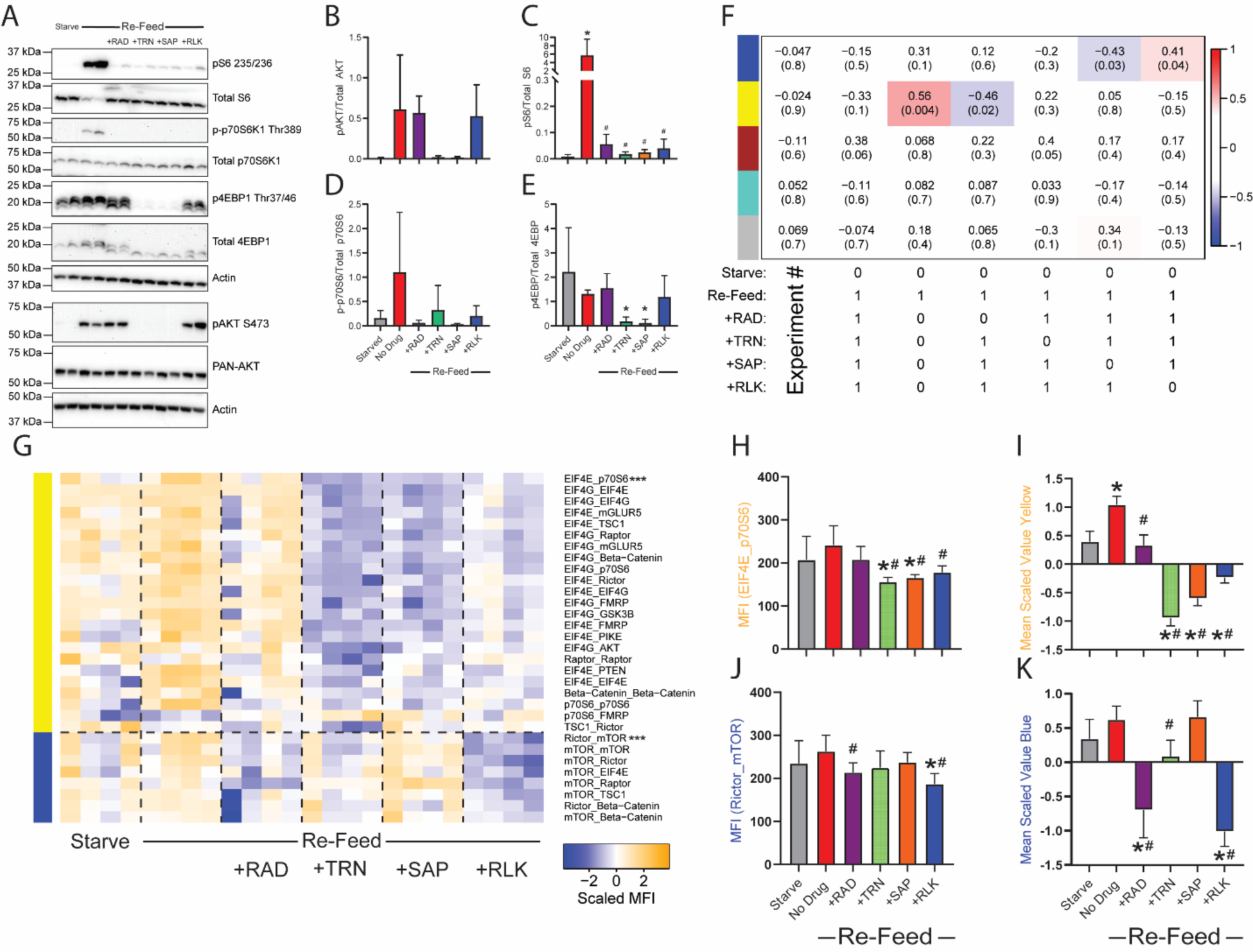
Different generations of mTOR Inhibitors differentially affect mTOR modules. A) Western blots of phospho- and total AKT, S6, p70S6K, 4EBP1 and actin for loading control, in 3T3 fibroblasts that were starved and re-fed in the presence of drugs. B-E) Quantification of blots in A. * indicates significantly different from starved, # indicates significantly different from re-fed by ANOVA followed by Bonferroni-corrected post-hoc testing, p<0.05. N=4. F) Module-trait table showing the correlation (top number) and the p-value (bottom number) between the eigenvector of each color-coded module (colored rectangles at left) and binary-coded trait labels (“hypotheses”) shown in the table below. G) Heatmap of the median scaled values of all significant interactions. Each box represents a single interaction measurement from a single biological replicate; columns correspond to a biological replicate while rows correspond to an interaction. Interactions are listed in descending order by module membership, color-coded by rectangles at left and by text color. N = 4. H) Median fluorescent intensity (MFI) of the interaction between EIF4E_p70S6, the interaction most correlated to yellow module behavior. I) Mean scaled value of all interactions in the yellow module, which corresponds to overall module behavior. J-K) Similar to H-I, but the blue modules. For H-K, * indicates significantly different from starved, # significantly different from re-fed, by ANOVA followed by Bonferroni-corrected post-hoc testing, p<0.05.

The blue module was composted of mTOR complex interactions similar to those observed in the blue module in Figure 1 (which responded to RAD but not to BKM), including mTOR_Raptor and mTOR_Rictor. These interactions were strongly suppressed by both RAD and RLK, but only weakly suppressed by TRN and not altered by SAP (Fig 4J,K). These data reflect a key difference in the mechanism of action between RAD-derived and ATP-competitive inhibitors: the former bind to FKBP12 and physically disrupt mTOR substrate accessibility (Yang et al., 2013), while latter bind the ATP binding site and inhibit mTOR in the absence of protein complex disruption.

### Hyperactivating mutations drive mTOR PIN activation

As a central regulator of cellular growth, the mTOR pathway is implicated in several human overgrowth disorders, including Tuberous Sclerosis, *PTEN* hamartoma tumor syndrome, Megalencephaly-capillary malformation syndrome, Focal Cortical Dysplasia, and hemimegaencephaly. Among the common pathogenic variants associated with these neurodevelopmental disorders, sometimes referred to as ‘mTORopathies” (reviewed in Karalis and Bateup, 2021), are mutations of *PIK3CA* or *MTOR*. The *PIK3CA* p.H1047R variant is the most common hyperactivating mutation in the catalytic p110 subunit of the PI3K holoenzyme that renders it resistant to inhibition by the p85 regulatory subunit, increasing basal PI3K activity (D’Gama et al., 2015; Gymnopoulos et al., 2007). The *MTOR* p.T1977I variant is a highly recurrent hyperactivating mutation in the FAT domain of mTOR, adjacent to the kinase domain, which may promote accessibility of substrates to the kinase domain (G. M. Mirzaa et al., 2016). We obtained fibroblasts from two individuals with overgrowth who had mutations in *PIK3CA*^H1047R^ (1y/o male) and *MTOR*^T1977I^ (13y/o female), as well as two unrelated healthy controls (4 y/o male and 9y/o female). While the cells were taken from genetically mosaic individuals, the mutant cells have an increased growth rate compared to wildtype cells (Mirzaa et al., 2016; Samuels et al., 2005), so the variable (mutant) allele frequency, measured by digital droplet PCR, was 49.9% for the PIKC3A mutant and 39.7% for the *MTOR* mutant, indicating 100% and 80% heterozygous mutant cells, respectively (Figure S7). Cells were serum starved for twelve hours and lysed immediately or stimulated with fresh serum-containing media for one hour before lysis. Western blot showed that the PIK3CA mutant fibroblasts displayed hyperphosphorylation of AKT and S6 during starvation (Fig 5A-C), while the mTOR mutant fibroblasts did not display any significant deficits in phosphorylation, consistent with previous reports (Di Donato et al., 2016; Grabiner et al., 2014).

**Figure 5:**
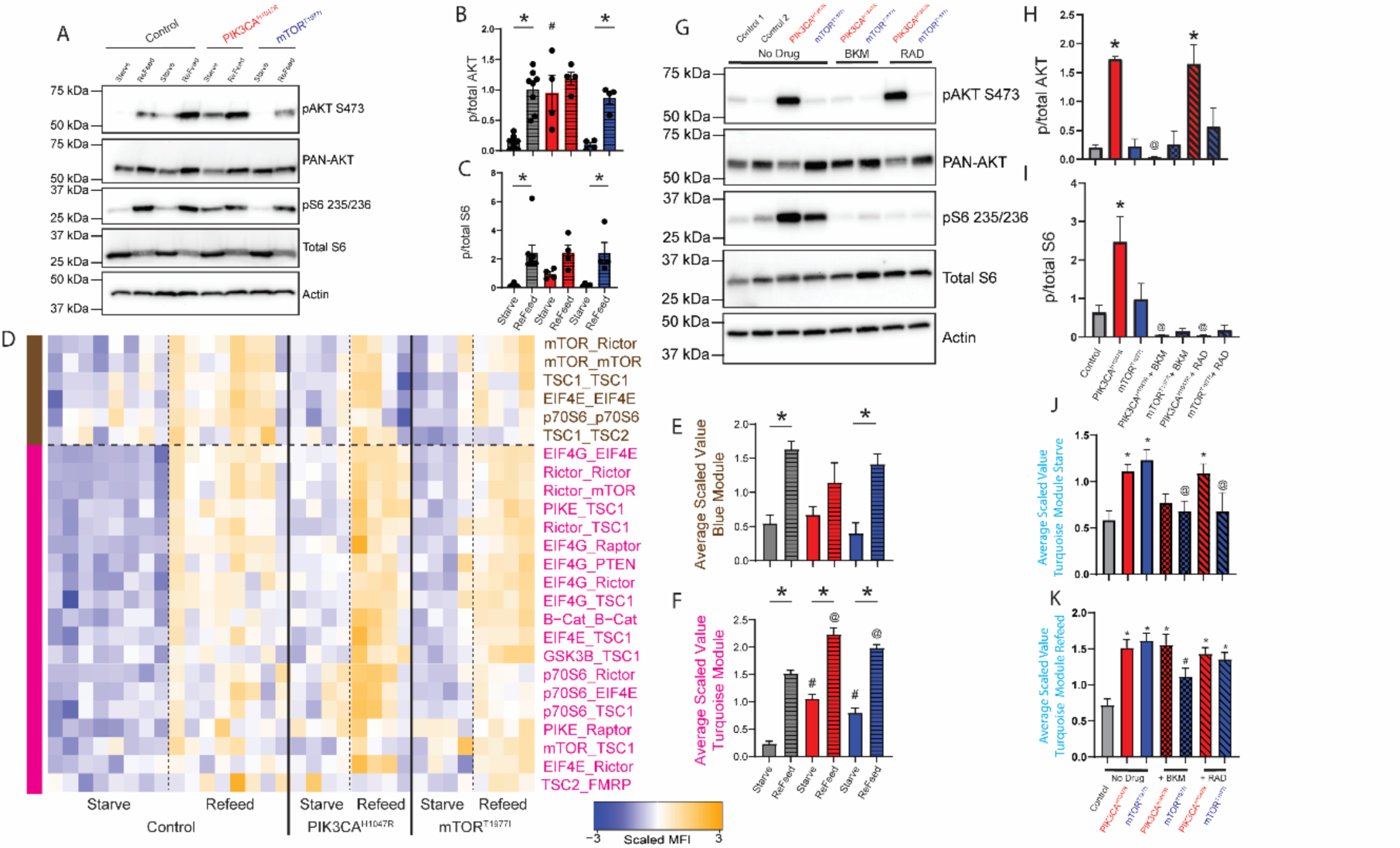
Activating mutations rescued by targeted drugs. Patient derived human fibroblasts with *PIK3CA^H1047R^* and *mTOR^T1977I^*gain of function mutations underwent a starve-refeed treatment. A) Representative western blots of phospho- and total AKT and S6, and actin for loading control. B-C) Quantification of blots in B. * indicates significantly different from starved within-genotype, # indicates significantly different from control-starved, by ANOVA followed by Bonferroni-corrected post-hoc testing, p<0.05. D) Heatmap of the median scaled values of all significant interactions. Each box represents a single interaction measurement from a single experiment; columns correspond to an experiment while rows correspond to an interaction. N = 4-8. Interactions are ordered by module, represented by colored bars on the left, and coloring of text. E) Mean scaled value of all interactions in the brown module. F) Mean scaled value of all interactions in the pink module. For E-F, * indicates significantly different from starved within-genotype, # significantly different from control-starved, and @ significantly different from control-refed, by ANOVA followed by Bonferroni-corrected post-hoc testing, p<0.05. G) Following starvation, patient-derived fibroblasts were treated with BKM120 or RAD001 during the re-feed period. Representative western blots of starved samples phospho- and total AKT and S6, and actin for loading control. H-I) Quantification of blots in B. * indicates significantly different from control, @ indicates significantly different from no-drug condition within-genotype, by ANOVA followed by Bonferroni-corrected post-hoc testing, p<0.05. J) Mean scaled value of all interactions in the turquoise module for starved, drug-treated patient fibroblasts. * indicates significantly different from control, @ indicates significantly different from no-drug condition within-genotype, by ANOVA followed by Bonferroni-corrected post-hoc testing, p<0.05. K) Mean scaled value of all interactions in the turquoise module for re-fed, drug-treated patient fibroblasts, similar to Figure 6J. * indicates significantly different from control, # indicates significantly different from no-drug condition within-genotype, by ANOVA followed by Bonferroni-corrected post-hoc testing, p<0.05. N=4-8.

Following QMI, we used CNA to independently identify two modules that correlated with refeeding (Fig 5D). A brown module, exemplified by mTOR_Rictor, was significantly elevated in re-fed control and *MTOR*^T1977I^ cells (compared to same-genotype starved cells), and trended towards significance in *PIK3CA*^H1047R^ cells (Fig 5E). There were no differences between genotypes in the starved or refed conditions. In contrast, a pink module, exemplified by EIF4G_EIF4E, was elevated in all comparisons tested (Fig 5F); not only did re-feeding increase module intensity compared to unstarved cells within-genotype, but comparisons of starved or re-fed mutant cells to starved or refed control cells were significantly elevated. PIK3CA^H1047R^ fibroblasts showed greater elevation than did MTOR^T1977I^ fibroblasts. These data are consistent with clinical data suggesting that strongly activating variants in PIK3CA are associated with more severe overgrowth than MTOR (Pirozzi et al., 2022). Specifically, the PIK3CA H1047R variant has been associated with severe somatic or brain overgrowth phenotypes including focal cortical dysplasia with severe epilepsy (G. Mirzaa et al., 2016), whereas the MTOR T1977I variant is associated with megalencephaly and polymicrogyria and overall less severe epilepsy (G. M. Mirzaa et al., 2016). Notably, our analysis did not identify additional interaction modules that were associated with hyperactivating mutations independently of re-feeding, indicating that nutrient status and hyperactivating mutations affected a similar set of protein complexes.

### Drug treatment of mutant fibroblasts

Most therapeutic approaches utilized for the ‘mTORopathies” focus on Rapamycin and its analogues, regardless of the location of the variant in the linear mTOR cascade (Karalis and Bateup, 2021; Nguyen and Bordey, 2021). However, given our results demonstrating differential regulation of the mTOR PIN by drugs targeting mTOR vs PI3K, we reasoned that a more targeted approach may improve network-scale rescue. We treated control, PIK3CA^H1047R^ and mTOR^T1977I^ fibroblasts with BKM or RAD to inhibit PI3K or mTORC1, respectively. Western blots from the starved samples for pAKT revealed control and mTOR^T1977I^ fibroblasts had low levels of p-AKT, while PIK3CA^H1047R^ had elevated basal p-AKT. BKM normalized pAKT in PIK3CA^H1047R^ fibroblasts, while RAD did not affect pAKT levels (Fig 5G-H). PIK3CA^H1047R^ and mTOR^T1977I^ fibroblasts showed elevated S6 phosphorylation, which was reduced following BKM and RAD treatment (Fig 5I). QMI, performed with a smaller antibody panel due to technical limitations, identified a turquoise module, exemplified by EIF4G_EIF4E, which was significantly elevated in both starved and re-fed PIK3CA^H1047R^ fibroblasts (Fig S8A,B). We graphed the average scaled value of the turquoise module for starved (Fig 5J) or re-fed cells (Fig 5K). For the re-fed condition, all mutant lines were hyperactive compared to control, and both mTOR-modifying drugs were ineffective, except for *mTOR*^T1977I^ fibroblasts treated with BKM (Fig 5K). However, for starved cells, we observed a significant increase in module activation in mTOR^T1977I^ and PIK3CA^H1047R^ fibroblasts compared to controls (Fig 5J). This increase was normalized by both BKM and RAD treatment in *mTOR*^T1977I^ fibroblasts, and by BKM treatment in *PIK3CA*^H1047R^ fibroblasts; however, RAD did not rescue hyperactivation in *PIK3CA*^H1047R^ fibroblasts (Fig 5J).

## Discussion

We developed a QMI panel targeted to key protein nodes in the mTOR pathway that allows direct measurement of mTOR PIN dynamics during signaling events, and we defined a PIN that is acutely modified by mTOR signaling. Following re-feeding with serum-derived growth factors, we observed a single module of coordinated protein-protein interactions that gradually increased over the course of an hour. Drugs targeting different nodes of the mTOR network, or drugs targeting mTOR using different mechanisms, altered complex sets of interactions (i.e. modules) that were not predicted by a linear pathway model. QMI offers an alternative way to characterize the mTOR signal transduction network, traditionally observed via changes in phosphorylation events, and allows access to a new level of biological complexity in the dynamic protein interactome.

### Kinetics of the mTOR PIN

In previous studies using QMI to monitor the T cell receptor signalosome and the glutamatergic synapse interactome (Lautz et al., 2021; Smith et al., 2016), peak signalosome activation occurred ∼5 minutes after stimulus presentation. Similarly, data presented here (Fig 1B-D) and in the literature (Rahman and Haugh, 2017) suggest phosphorylation of AKT peaks ∼5 minutes after stimulus presentation, and downstream activation of S6 peaks ∼15 minutes. However, QMI observed virtually no PIN response at 5 minutes, and only a small, partial response at 15 minutes, with peak response not occurring for at least an hour. Additionally, the mTOR network still did not recover to its pre-starvation state even after an hour-long response. These data imply that changes in phosphorylation status are slow to translate into changes in protein complexes, which is unexpected because changes in the phosphorylation status of interacting proteins is thought to be directly responsible for changes in protein-protein interactions. For downstream interactions such as EIF4E_EIF4G, which represent the final transcriptional output of the mTOR system, this delay could be explained by a slow rate of signal progression through a linear cascade. However, for more upstream interactions such as RICTOR_mTOR (representing MTORC2) or PIKE_TSC1, the reason for the slow kinetics is unclear. Future work to reconcile the slow rate of network modification with the much faster rate of phosphorylation is warranted.

### Linear vs. Network models of drug specificity

By adding different small molecule inhibitors of mTOR pathway nodes, we were able to fracture the single module of co-regulated interactions that responded to refeeding into two main modules of interactions, blue and yellow, that responded differently to each type of inhibition. We summarize the modules that we identified throughout this study in Figure 6. We expected that the modular organization of the network would follow the traditional linear model, with BKM inhibiting all modules, RAD inhibiting the fewest number of modules, and AZD being intermediate. Moreover, we expected Rapalogs and next-generation mTOR inhibitors to produce qualitatively similar changes to interaction modules, even if the magnitude of inhibition was different. Surprisingly, the modules did not follow this pattern.

**Figure 6:**
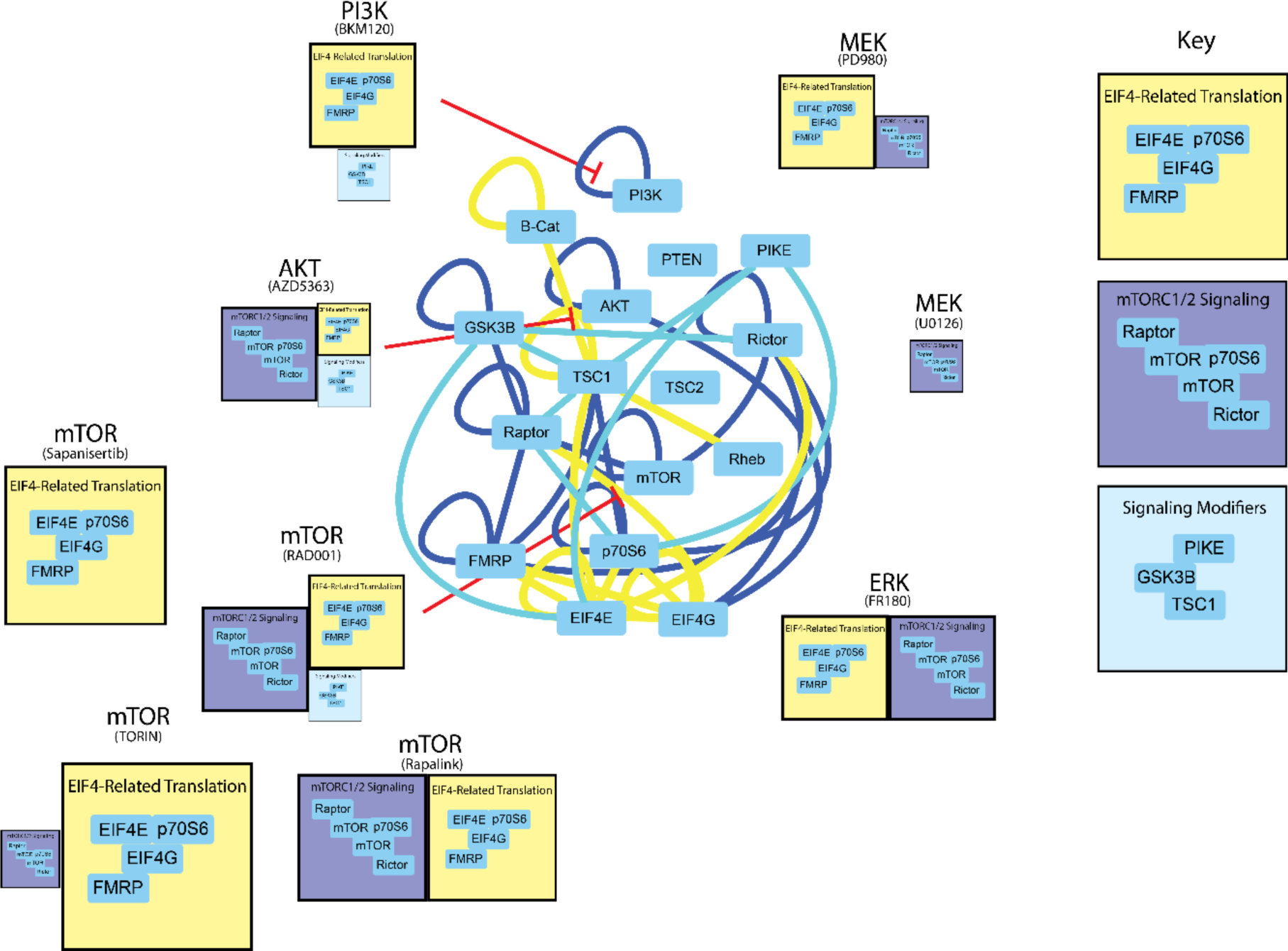
Modular organization of the mTOR PIN. Node-edge diagram represents all significant interactions that changed with re-feeding during the mTOR and ERK inhibitor experiments (Fig 2, 3, and 4). Edge color indicates the assigned module of each interaction. The boxes below each drug target indicate the modules that were inhibited by the drug; the size of the box corresponds to the intensity of inhibition, and the content of all similarly-colored boxes is consistent throughout.

The yellow “downstream” module was inhibited by all drugs except U01, and most strongly by BKM and the next generation mTOR inhibitors. It is somewhat counterintuitive that BKM produced stronger inhibition than RAD in head-to-head studies, since the yellow module contains interactions among the most downstream proteins (EIF4E and G, p70S6, and FMRP). Given that phosphorylation of p70S6 is mediated by both mTOR and PDK1 (Pullen et al., 1998), it is possible that BKM, by inhibiting PDK1 downstream of PI3K, provides double-inhibition not seen in AZD and RAD. It is also possible that ATP-competitive inhibitors like BKM, SAP and TOR provide more complete mTOR inhibition than RAD. Indeed, next-generation inhibitors are known to more completely inhibit mTORC1 and mTORC2 compared to rapamycin derivatives (as evidenced by 4EBP1 and AKT phosphorylation), and are more effective at inhibiting the growth of cancer cells *in vitro* and in mouse models (Gökmen-Polar et al., 2012; Liu et al., 2010; Rodrik-Outmezguine et al., 2016). Moreover, it has been suggested some mTOR signaling is RAD-resistant, but may be inhibited amino acid starvation (Peng et al., 2002), perhaps leading to a network state that resembles our next-generation mTOR inhibitor data. Future work comparing amino acid starvation to next-generation mTOR signaling may be warranted. Overall, these data suggest that the yellow module is a core mTOR-dependent readout, that is more effectively inhibited by ATP-competitive kinase inhibition of PI3K or mTOR than by Rapalogs.

The blue module is more difficult to interpret. This module was inhibited by RAD, AZD and ERK inhibition, and weakly inhibited by MEK inhibition, which suggests that mTOR signaling downstream of both AKT and ERK may mediate module behavior. However, PI3K inhibition and mTOR kinase inhibition by SAP were completely ineffective at preventing blue module activation, and mTOR kinase inhibition by TRN was weak, especially compared to the magnitude of yellow module inhibition. Moreover, the ratio of inhibition of the blue module compared to the yellow module was different for each drug; for example ERK inhibition with FR1 produced equal inhibition of each module, AKT inhibition with AZD produced stronger inhibition of the blue module, and mTOR inhibition with TRN produced stronger inhibition of the yellow module.

Perhaps these differences reflect uncharacterized off-target effects on other relevant kinases, or are due to uncharacterized feedback loops or other non-linear relationships among the mTOR network. For example, inhibiting MEK with U01 vs PD9 produced strikingly different patterns of PIN inhibition. PD9 was less effective at inhibiting ERK phosphorylation by western blot compared to U01, but was much more effective at inhibiting “yellow” module interactions containing EIF4E/G and FMRP. Despite sharing a mutually exclusive binding site on MEK, and despite U0126 binding with a 100-fold higher affinity to MEK (Favata et al., 1998), U0126 failed to inhibit the yellow module. While both PD98059 and U0126 have known off-target (i.e. non-MEK-mediated) effects, for example, activating AMP-activated protein kinase (Dokladda et al., 2005) or Kinase Suppressor of RAS (KSR-1) (Wang and Studzinski, 2001), these off-target effects are thought to be similar for both drugs. Similarly, the next-generation mTOR inhibitor experiments identified clear differences in “blue” module inhibition between TRN and SAP, which also function through similar ATP-competitive mechanisms. These data are reminiscent of a recent phospho-mass spectrometry study comparing 5 different AKT inhibitors, which showed only limited overlap: of 1700 altered phospho-peptides, only 276 were perturbed by all 5 compounds (Wiechmann 2021). The strikingly different effects of the inhibitor drugs on the mTOR PIN highlights the emerging concept that “specific” inhibitor drugs are often not very specific at all. A potential advantage of QMI-based network measurements is that they can highlight differences between inhibitor drugs better than traditional phospho-western readouts.

### Correcting pathogenic network hyperactivity

The human patient fibroblast model gives us the ability to observe how an overactive member of the mTOR cascade contributes to the response of the PIN. Hyperactivity at the top of the mTOR pathway, at PI3K, produced more robust differences than further downstream at mTOR, consistent with our inhibitor data, as well as with clinical data showing more severe phenotypes in patients with PI3K vs mTOR mutations (G. Mirzaa et al., 2016). PI3K and mTOR mutations did not produce novel PIN states; rather, they increased the fluorescent intensity of the interactions that composed the stimulation-responsive module in both the starved and re-fed states compared to wildtype. Importantly, we did not identify modules that correlated with the presence of a mutation, independent of starvation/refeeding, which would have indicated changes to the mTOR network independent of nutrient sensing. These data imply that simple inhibition of signaling may normalize the dynamic range of the system. Indeed, BKM was able to normalize PI3K hyperactivation, and RAD and BKM were both able to normalize mTOR hyperactivity. However, RAD was not able to normalize the hyperactive network state produced by PI3K mutation, likely because PI3K has direct links to downstream signaling nodes that bypass mTOR, as demonstrated also by the BKM inhibition data. Future experiments should focus on how mTOR inhibitors might affect mTOR network hyperactivation at more physiologically relevant states, outside of the context of serum-starvation, since the effect of hyperactivation in non-physiological starvation conditions that may not be fully applicable to an untreated condition.

This result, while somewhat intuitive, speaks directly to clinical trials occurring in overgrowth disorders, cancer, or ASD-related mTOR-opathies. Ongoing translational research is attempting to correct upstream mutations in PI3K (Byeon et al., 2020; Parker et al., 2019), PTEN (Schmid et al., 2014), or TSC (Krueger et al., 2017; Overwater et al., 2019) with rapamycin or its analogs, with mixed outcomes. Perhaps this treatment strategy, based on linear models implying upstream effects filter through mTORC1, ignores the complexity of the pathway. Our data, as well as recent work correcting PI3K-dependent (Roy et al., 2021) or TSC2-dependent (Nguyen et al., 2022) epilepsies with more targeted drugs, imply that direct correction of a hyperactive node may be more effective than the simple use of rapalogs for all mTOR pathway disorders.

### Limitations

A general limitation of QMI is that it measures the epitope-specific binding of commercially available antibodies, so differences in fluorescent signal may be due to differences in the targeted protein-protein interaction, or due occlusion of an antibody binding site by an unknown protein joining a complex. While we validated some interactions by IP-western blotting, the sensitivity of QMI exceeds that of IP-westerns (Smith et al., 2012). Thus, while the overall behavior of modules observed by QMI is robust, the behavior of any specific interaction may require further validation.

A second limitation of this study is that the inhibitors used to block specific kinases may have affected other kinases whose inhibition may have contributed to the observed changes in the mTOR PIN. With the exception of the next-generation mTOR inhibitor experiments, we used single concentrations of drugs based on previously published reports, so we cannot rule out off-target effects. Future work using genetic knockouts or knockdown of mTOR components could address these limitations, but also presents further difficulties due to the requirement of mTOR signaling for growth and cell cycle, as well as due to extensive feedback loops within the mTOR network.

Finally, we focused on a single starve-refeed paradigm, while other avenues exist to either activate (i.e. growth factor stimulation) or inhibit (i.e. amino acid starvation) the network. Future experiments using QMI to more fully characterize the mTOR signal transduction network could further disentangle the complex relationships between experimental manipulations and the response of the mTOR signal transduction network.

## Materials and Methods

### Cell Culture

NIH 3T3 mouse fibroblasts were acquired from the American Type Culture Collection. The cells were cultured at 37° C at 5% CO2 in Dulbecco’s modified Eagle’s medium supplemented with 10% fetal bovine serum, 1 % non-essential amino acids [Gibco-11140],1% GlutaMax [Gibco-35050], and 1 % penicillin and streptomycin.

Patient derived fibroblast were obtained from punch biopsies of affected children enrolled in the IRB-approved developmental brain disorders research program. Fibroblasts were cultured at 37° C at 5% CO2 in Dulbecco’s modified Eagle’s medium with nutrient mixture F-12 (Gibco-11330) and supplemented with 15% fetal bovine serum, 1 % non-essential amino acids [Gibco-11140],1% GlutaMax [Gibco-35050], and 1 % penicillin and streptomycin. ddPCR was performed on validated probes using a QX200 ddPCR system following the manufacturer’s instructions, as detailed in (Pirozzi et al., 2022).

### Antibodies and Other Reagents

The following antibodies for the purpose of western blotting were acquired from Cell Signaling Technologies: pAKT S473 (Catalog number 4060), PAN-AKT (#2920), Phospho-p44/42 MAPK (#4370), p44/42 MAPK (#4695), S6 Ribosomal Protein (#2317), Phospho-S6 Ribosomal Protein (#4858), Phospho-p70S6 Kinase Thr389 (#97596), p70S6 Kinase (#9202), Phospho-4EBP1 Thr37/46 (#2855), 4EBP1 (#9452). Antibodies against Beta-Actin were purchased from GeneTex (catalog number 109639). MAPK inhibitors were purchased from Tocris: FR180204 (catalog number 3706), U0126 (catalog number 1144), PD980595 (catalog number 1213). Sapanisertib (Cat# HY-13328) and Rapalink-1 (Cat# HY-111373) were purchased from MedChem Express. Torin-1 (Cat #4247) was purchased from Tocris. PI3K/AKT/mTOR inhibitors BKM120 (Buparlisib; Novartis, Switzerland), AZD5363 (Capivasertib; Selleckchem, USA), and RAD001 (Everolimus, Chem Express Cat# 159351-69-6) were generously provided by Kathleen Millen.

### Serum Starvation, Stimulation, Drug Treatment, and Lysate Preparation

Fibroblast cultures were grown to ∼70-80% confluence in 10-cm dishes or 6-well plates and were serum starved for 12 hours. During pharmacological inhibition experiments, inhibitors were added at the eleventh hour of starvation. After starvation, cells were re-fed with fresh media. Inhibitors were present during re-feeding in the respective experiments. Inhibitors were used at the following concentrations: BKM120 (2 uM), AZD5363 (1 uM), RAD001 (40 nM), FR180204 (100 uM), U0126 (25 uM), PD980595 (50 uM), TORIN1 (40 nM), Sapanisertib (125 nM), and Rapalink-1 (10 nM). Media was removed, and cells were washed in PBS and scraped on ice with ice cold lysis buffer [1% Digitonin, 150 mM NaCl, 50 mM Tris (pH 7.4), 10 mM NaF, 2 mM sodium orthovanadate, protease inhibitor cocktail (Sigma-Aldrich), and phosphatase inhibitor cocktail (Sigma-Aldrich)] and transferred to a centrifuge tube. After fifteen minutes of incubation on ice, samples were centrifuged at high speed for fifteen minutes to remove nuclei and debris. The protein concentration of the lysate was determined with a Bradford assay (Pierce).

### Quantitative Multiplex Immunoprecipitation

QMI was performed as described previously (Brown et al., 2019). A master mix containing each antibody-coupled Luminex bead was prepared and distributed to lysates normalized for protein concentration. Samples were incubated overnight at 4°C on a rotator. The following day, samples were washed in cold FlyP buffer [50 mM tris (pH7.4), 100 mM NaCl, 1% bovine serum albumin, and 0.02% sodium azide] and distributed into twice as many wells of a 96-well plate as there were probe antibodies for technical replicates. Biotinylated probe antibodies were added and the plate was incubated at 4°C with gentle agitation for one hour. The resulting complexes were washed three times with FlyP buffer on an automatic plate washer. The samples were then incubated for thirty minutes with streptavidin-phycoerythrin at 4°C with gentle agitation. Samples were washed three times again and resuspended in 120 ul of cold FlyP buffer and processed with a customized refrigerated Bio-Plex 200.

### Adaptive nonparametric analysis with an empirical alpha cutoff (ANC)

Statistically significant differences in bead distributions between conditions for each of ∼400 individual interactions, after correcting for multiple comparisons, were identified using ANC as described in previous work (Brown et al., 2019; Smith et al., 2016). Any Interaction that was found to be significant by an ANC comparison was considered a “hit.”

### Correlation Network Analysis (CNA)

Modules of Interactions that covaried with experimental conditions were identified using weighted correlation network analysis (Langfelder and Horvath, 2008) as described in previous work (Brown et al., 2019; Smith et al., 2016). Bead distributions used in ANC were collapsed and the median fluorescent intensity (MFI) value was averaged across technical replicates for input into the WGCNA package for R. Interactions with an MFI less than 100 were removed as noise, and batch effects were corrected using COMBAT (Leek et al., 2012). Power values giving the approximation of scale-free topology were determined using soft thresholding with a power adjacency function, and modules were determined by the TOM matrix function in WGCNA. Modules whose eigenvectors were correlated with an experimental trait (P < 0.05) were of interest. Interactions whose probability of membership in a module of interest was (P < 0.05) were considered “hits”. Interaction that were “hits” by both ANC and CNA for a given experimental condition were considered high confidence interactions affected in that condition.

### Hierarchical Clustering and PCA

Post-COMBAT, log_2_ transformed MFI values were clustered using the hclust function in R with a correlation distance matrix and average clustering method. Approximately unbiased (AU) P values were determined using the pvclust package in R. PCA was performed using the prcomp function in R.

## Acknowledgements

We thank the people who generously donated knockout cell lines or lysates for use as negative antibody controls: Andrew Hsieh, David Sabatini, David Kwiatkowski, Paul Titchenell and Paul Monga. We would also like to thank Kathleen Millen for providing reagents and insight, including editing the manuscript, Filomena Pirozzi for sharing insights regarding experimental design, protocols and troubleshooting of patient-derived fibroblast experiments, and Smita Yadav for thoughtful discussions. This work was supported by grants MH103545 (SEPS), Jordan’s Guardian Angels (G.M.M), the Sunderland Foundation (G.M.M) and the Brotman Baty Institute (G.M.M. and D.T.W). Special gratitude to Barbara and Eric Mann for their support of our research (G.M.M.), as well as well as all donors to Seattle Children’s Research Institute who invest in breakthrough discoveries that help prevent, treat, and eliminate childhood disease. The authors declare no conflicts of interest.

## Author Contributions

DTW and SEPS designed the study. DTW and CSB performed all experiments. JS characterized the patient-derived mutant fibroblasts. GMM obtained the fibroblasts and assisted with the design and analysis of all experiments performed on patient-derived cell lines. DTW, JS, GMM and SEPS analyzed data. DTW and SEPS wrote the manuscript. All authors read and approved the manuscript.

## Notes

### Competing Interest Statement

The authors have declared no competing interest.

