## Supplemental Figures and Tables for "Protein interaction network analysis of mTOR signaling reveals modular organization"

**for:**

### Multiplex network analysis of protein interactions mediating TOR signal transduction

**Supplement contains:**

**Figures S1-S7**

**Table S1**

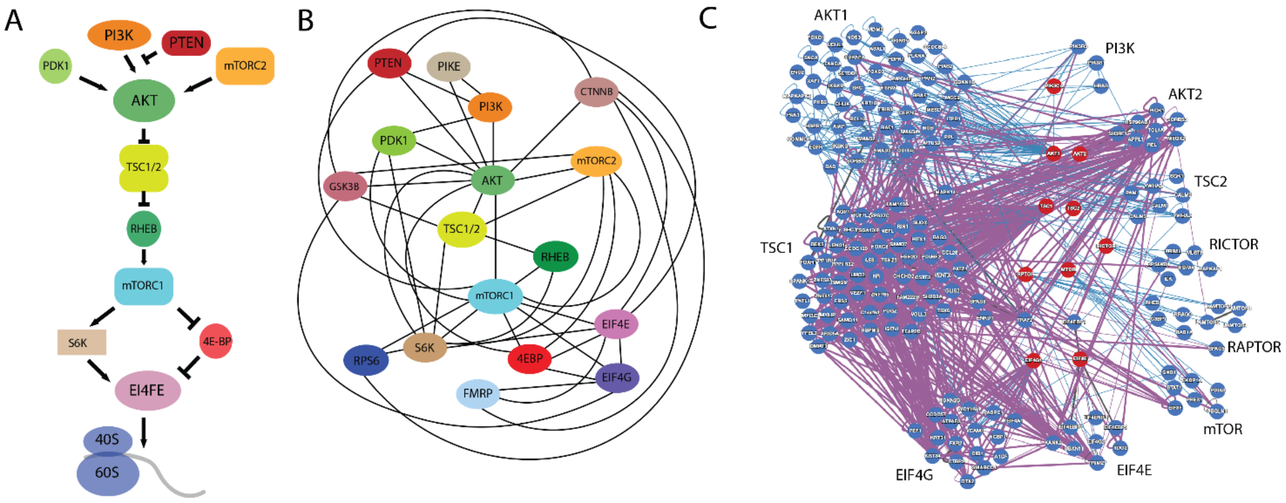


**Figure S1: Linear vs. Network modeling of mTOR signal transduction.** A) A linear model highlights how each node of the mTOR network affects the next in a simple, organized "cascade". Arrows indicate activation, lines indicate inhibition. B) Known protein-protein interactions in human or mouse cell lines among members of the mTOR signal transduction network, as listed in BioGrid and String databases. The nodes shown are the proteins selected for inclusion in the mTOR QMI panel, lines indicate a documented physical interaction. C) Protein-protein interactions in the Human Reference Interactome database (Luck et al., 2020), based on yeast 2-hybrid screening in immortalized human cells. Red nodes are queried members of the mTOR linear pathway, blue nodes are first-degree interactors, and edges indicate all documented binary interactions between the protein nodes shown. Red nodes are arranged by pathway hierarchy, blue nodes are clustered by interaction partner as labeled.

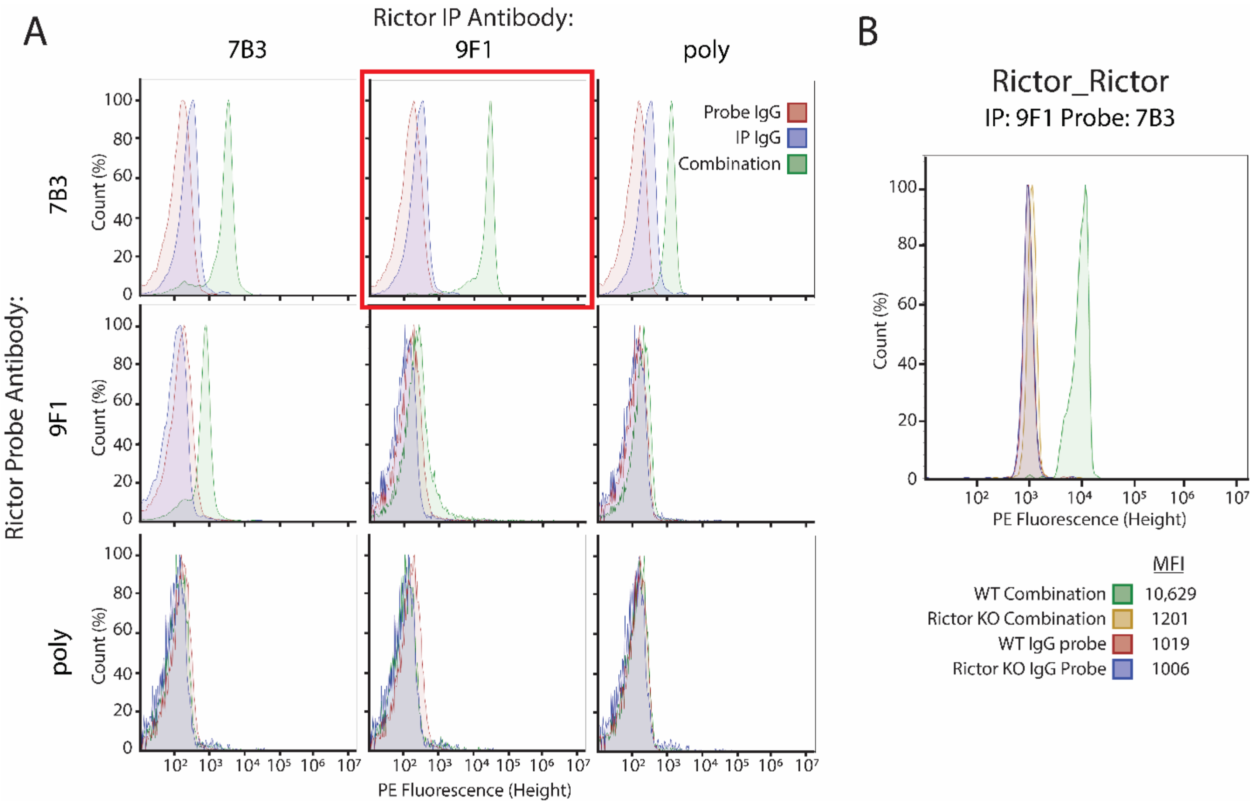


**Figure S2. Specificity validation of QMI antibody pairs in mouse.** A) Several candidate antibodies were bead-coupled (columns) and biotinylated (rows), and all IP-probe combinations were tested. Histograms for each IP_probe pair (green) were compared to IgG_probe (red) and IP_IgG (blue) controls, and the pair that produced the strongest signal over background was selected (7B3_9F1). B) Specificity was confirmed by comparing wildtype brain lysate to brain lysate from a KO mouse (yellow), which overlapped with IgG controls (red, blue).

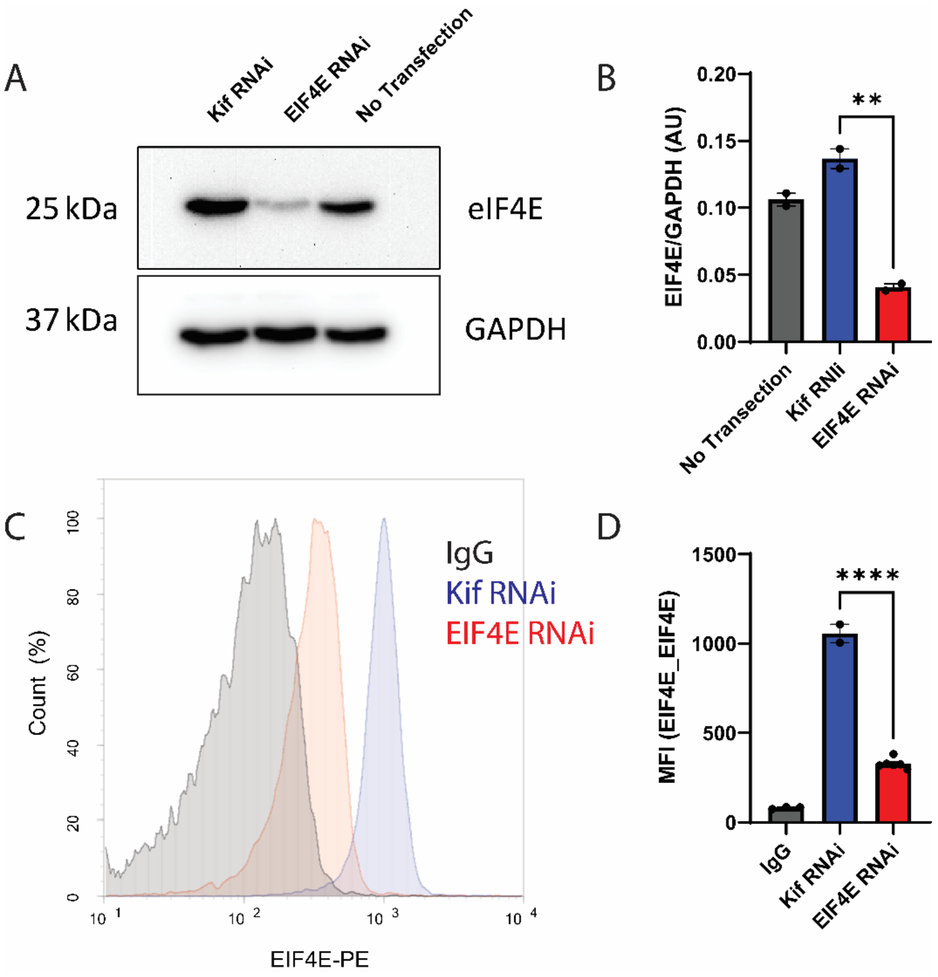


**Figure S3: Specificity validation of QMI antibody pairs in human** (A) Western blots from HEK 293 cells treated with RNAi to knockdown EIF4E. (B) Quantification of A, normalized to GAPDH levels. ** indicates significantly different from the KIF positive control by ANOVA followed by Bonferroni-corrected post-hoc testing, $p<0.05$. (C) Human specificity was confirmed by comparing KIF treated 293 cells (blue) to EIF4E RNAi treated 293 cells (red) and IGG precipitated samples (grey). (D) Quantification of the median fluorescent values from C.

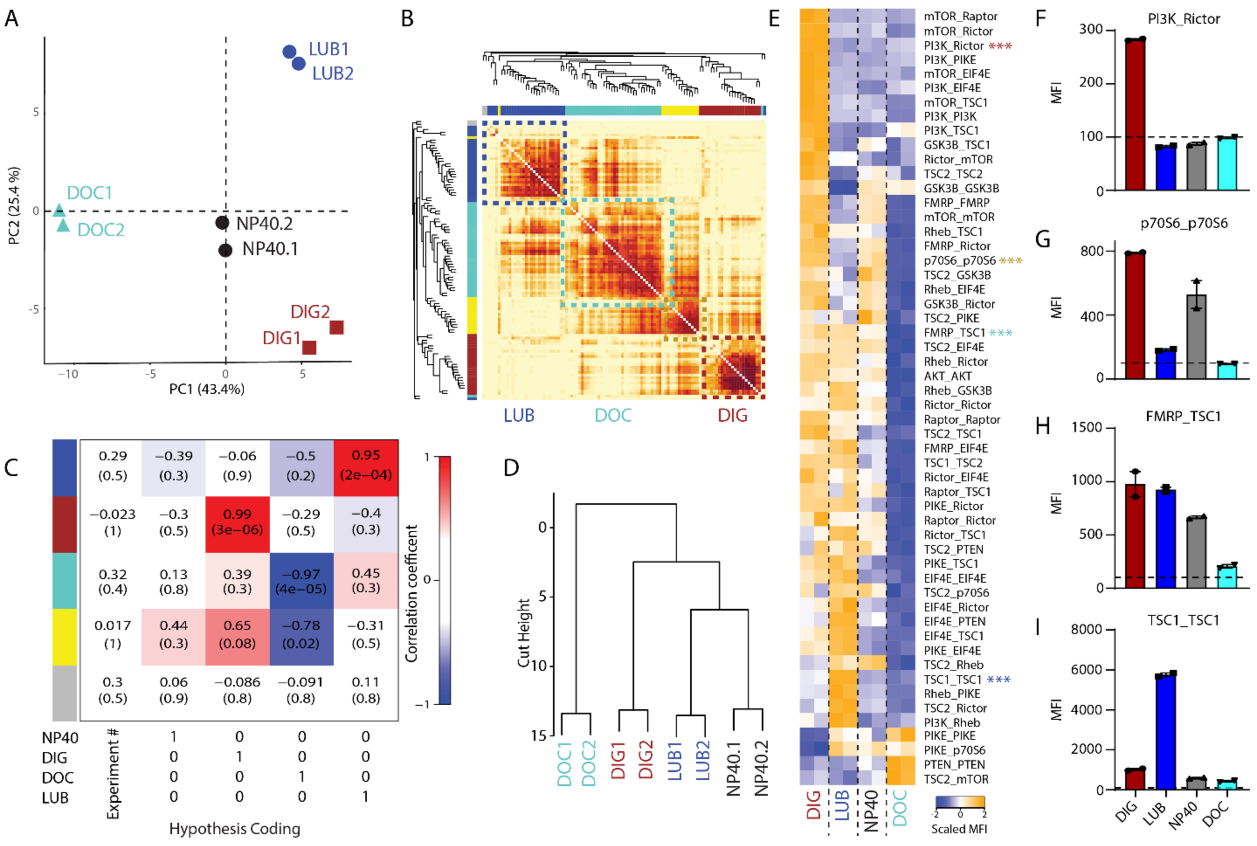


**Figure S4: Detergent affects protein complexes detected by QMI** (A) Principal component analysis (PCA) of P0 mouse cortical lysates lysed in four different detergents: Deoxycholate (DOC-turquoise), NP-40 (black), Lubrol (LUB-blue), and Digitonin (DIG-red) N=2 per condition. (B). A topological overlap matrix (TOM) reveals four modules of correlated PiSCES; dashed boxes outline each arbitrariliy-colored module. (C). Module-trait table showing the correlation (top number) and the p-value (bottom number) calculated by CNA for each trait. Hypothesis coding table below indicates the binary coding of detergent “traits”. (D). Hierarchical clustering of QMI data showing strong correlation between detergent repliates, and seperation of data by detergent  (E). Heatmap of the median scaled values of all dynamic interactions that showed significant changes between deteregent conditions. Each box represents a single interaction measurement from a single experimental replicate; columns correspond to a replicate while rows correspond to an interaction. N=2. (F-I). Bar graphs of the median fluorescent intensity (MFI) from a representative interaction in each module across samples.

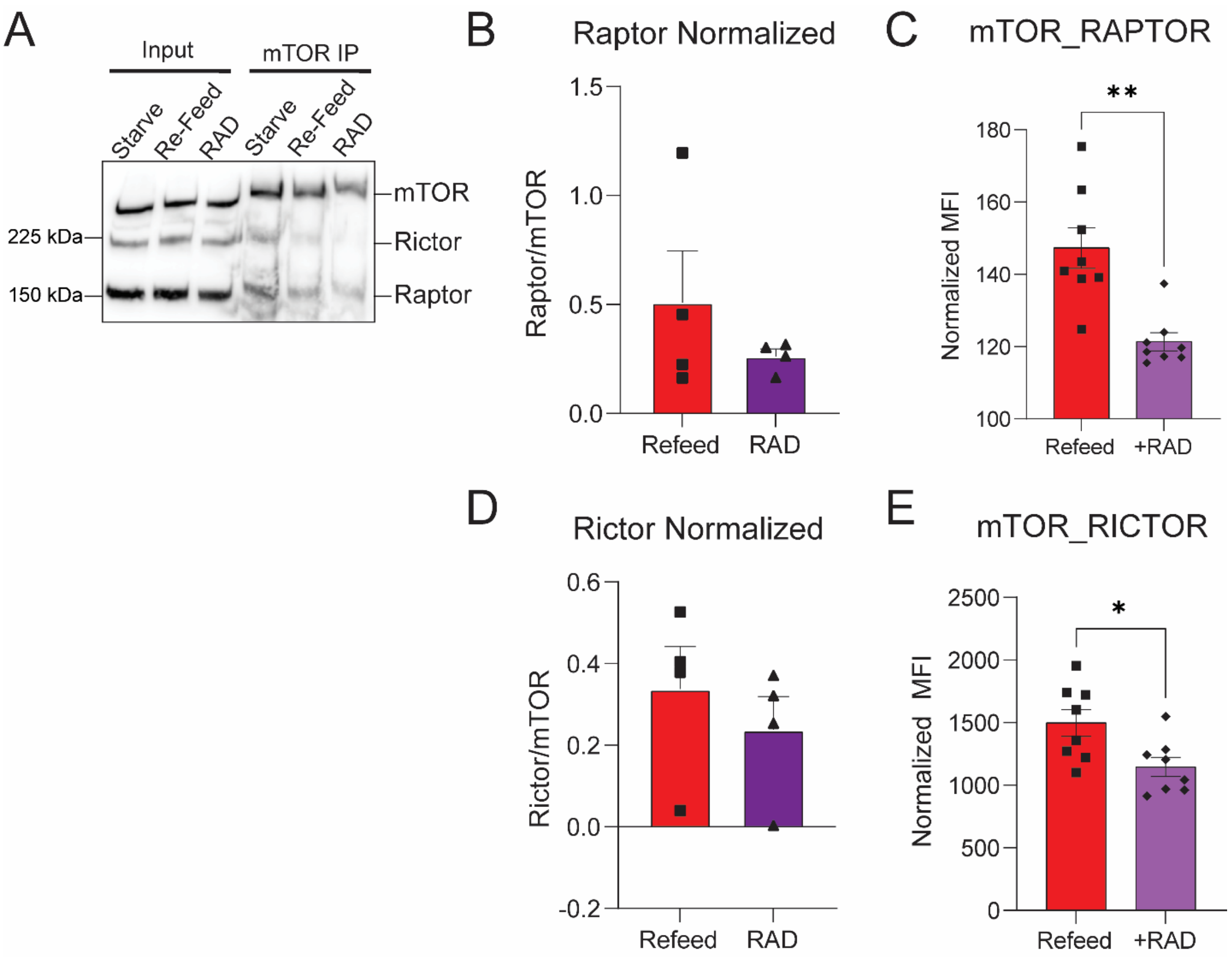


**Figure S5: Co-immunoprecipitation of mTOR following serum starvation and RAD001 treatment** (A) IP-western blots from 3T3 fibroblasts underwent serum starvation and refeeding protocol with or without 40 nM RAD001 and lysed in 0.3% Chaps buffer. (B) Quantification of Raptor probe from Refed and RAD treated samples normalized to mTOR. (C) The normalized median fluorescent values of the mTOR_RAPTOR interaction from Figure 2. (D) Quantification of Rictor probe from Refed and RAD treated samples normalized to mTOR. (C) The normalized median fluorescent values of the mTOR_RICTOR interaction from Figure 2.

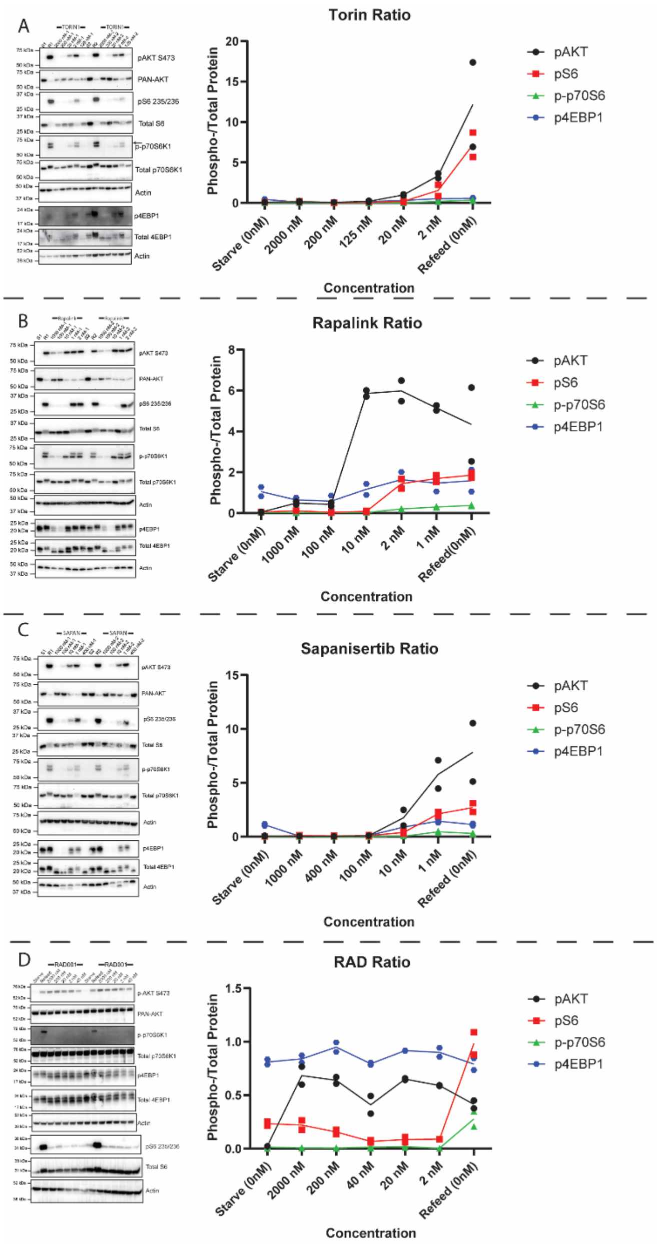


**Figure S5: Phosphorylation level curves related to Figure 4.** A) Phospho-western blots for AKT (black), S6 (red), p70S6 (green), and 4EBP1 (blue) from a serum-starvation experiment in the presence of various concentrations of TORIN1 along with a graph of each respective phosphorylation level at different concentrations. B-D) Similar figures as S4A but for Rapalink, Sapanisertib, and RAD001 respectively.

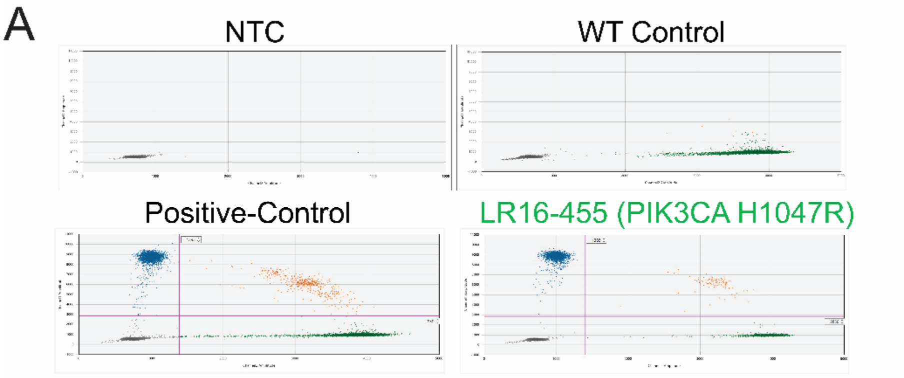


| Sample | Date | VAF (%) | VAF Max (%) | VAF Min (%) | Delta CI (%) | Accepted Droplets | Mut Positive | WT Positive | Probeset |
|---|---|---|---|---|---|---|---|---|---|
| NTC | 11/29/2021 | 0.00 | 0.00 | 0.00 | | 55721 | 0 | 1 | H1047R |
| WT Control | 11/29/2021 | 0.03 | 0.06 | 0.00 | | 52789 | 4 | 12633 | H1047R |
| Pos Con LR18-349 1702094 (22%) | 11/29/2021 | 24.99 | 25.78 | 24.21 | 24.15 | 57679 | 2956 | 8424 | H1047R |
| LR16-455 1702093 | 11/29/2021 | 49.94 | 51.29 | 48.60 | 48.54 | 64892 | 2661 | 2667 | H1047R |

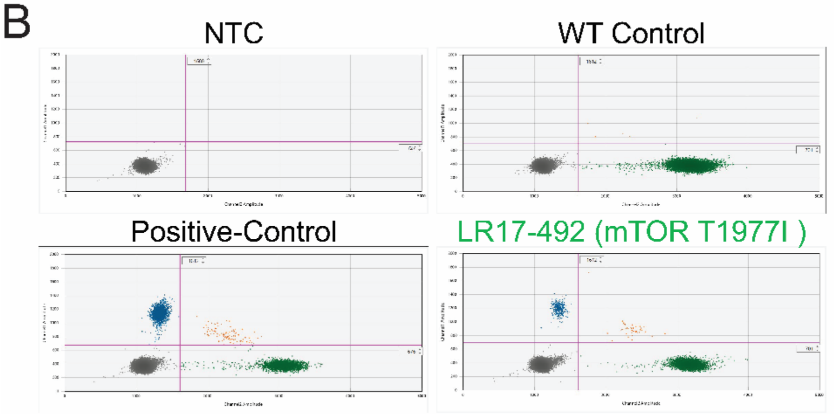


| Sample | Date | VAF (%) | VAF Max (%) | VAF Min (%) | Delta CI (%) | Accepted Droplets | Mut Positive | WT Positive | Probeset |
|---|---|---|---|---|---|---|---|---|---|
| NTC | 11/9/2021 | | | | | 57496 | 0 | 1 | T1977I |
| WT Control | 11/9/2021 | | | | | 61929 | 6 | 19565 | T1977I |
| LR17-492 1802021 Pos Con (41%) | 11/9/2021 | 39.32 | 40.66 | 37.98 | 37.98 | 63552 | 2016 | 3084 | T1977I |
| LR17 492 | 11/9/2021 | 5.15 | 5.66 | 4.65 | 4.65 | 58556 | 379 | 5598 | T1977I |

**Figure S6: Droplet digital PCR of patient derived mutant mTOR PIK3CA and mTOR** (A) Droplet digital PCR results for the PIK3CA H1047R line, summarized in a table. (B) Droplet digital PCR results for the mTOR T1977I line, summarized in a table.

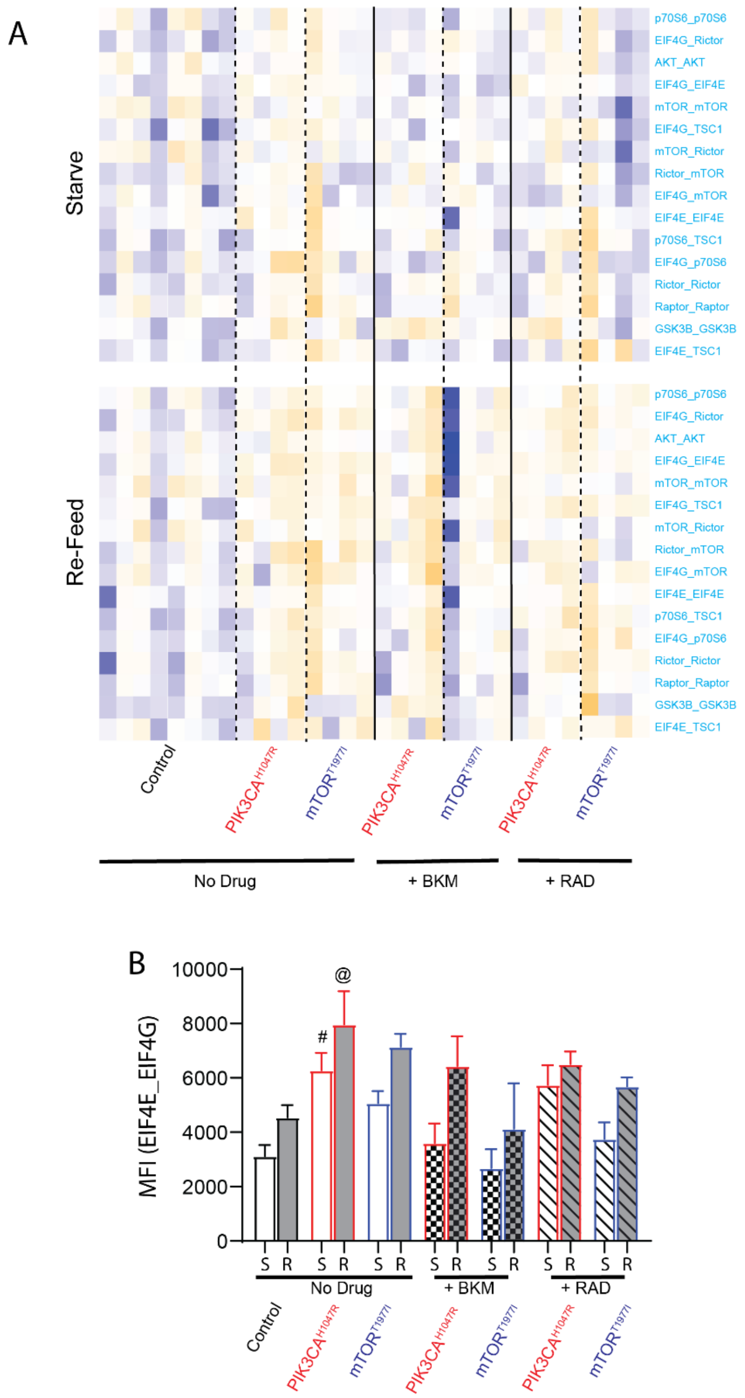


**Figure S7: Mutant fibroblast drug inhibition experiment, related to Figure 5.** A) Heatmap of the median scaled values of all dynamic PiSCES that showed ANC$\cap$CNA-significant changes. Each box represents a single interaction measurement from a single experimental replicate; columns correspond to an experimental replicate while rows correspond to an interaction. N = 4-8. B) Median fluorescent intensity (MFI) of EIF4E_EIF4G, the interaction most correlated with the turquoise module, shown for all conditions. * indicates significantly different from starved within-genotype, # significantly different from control-starved, and @ signficantly different from control-re-fed, by ANOVA followed by Bonferroni-corrected post-hoc testing, $p<0.05$. N=4-8

### Table S1: List of antibody clones and validations.

| | Target | Uniprot identifier | Uniprot # | IP antibody clone | IP antibody Catalog number | IP antibody Supplier | Probe antibody clone | Probe antibody Catalog number | Probe antibody Supplier | Mouse Validation | Human Validation |
|---|---|---|---|---|---|---|---|---|---|---|---|
| 1 | AKT1 | AKT1_MOUSE | P31750 | SKB1 | 05-591 | Millipore | B1 | sc-5298 | Santa Cruz | siRNA | RNAi |
| 2 | Beta-Catenin | CTNB1_MOUSE | Q02248 | 196624 | MAB13292 | R&D Systems | OTI2H7 | NBP2-03615 | Novus Biologicals | Mouse KO liver lysate (B6.129-Ctnnb1tm2Kem/KnwJ) | RNAi |
| 3 | eIF4E | IF4E_MOUSE | P63073 | 299910 | MAB3228 | R&D Systems | Poluclonal | A301-153A | Bethyl | siRNA | RNAi |
| 4 | EIF4G | IF4G1_MOUSE | Q6NZJ6 | G6899 | MA5-14971 | ThermoFisher-Scientific | 3A10 | SAB1403762 | Sigma-Aldrich | siRNA | |
| 5 | FMRP | FMR1_MOUSE | P35922 | Polyclonal | F4055 | Sigma-Aldrich | 5C2 | MMS-5232 | Biolegend | Mouse KO brain lystate (B6.129P2-Fmr1tm1Cgr/J) | RNAi |
| 6 | GSK3B | GSK3B_MOUSE | Q9WV60 | 3D6B4 | NBP2-52478 | Novus Biologicals | 3D10 | NBP1-47470 | Novus Biologicals | Mouse Embryonic fibroblast lysate (GSK3β(-/-); GSK3α(flox/-) (3/4KO) | RNAi |
| 7 | mGluR5 | GRM5_MOUSE | Q3UVX5 | 5675 | AB5675 | Millipore | N75/3 | 75-115 | Neuromab | Plasmid overexpression (Lautz et al 2018) | |
| 8 | mTOR | MTOR_MOUSE | Q9JLN9 | 2ID8.2 | 05-1592 | Millipore | 30 | sc-517464 | Santa Cruz | siRNA | RNAi |
| 9 | p70S6K | KS6A1_MOUSE | P18653 | 215247 | MAB8962 | R&D Systems | Polyclonal | AF8962 | Novus Biologicals | Mouse KO liver lysate (C.129(B6)-Rps6kb1tm1Gtho/NsoJ) | RNAi |
| 10 | PI3K | P85A_MOUSE | P26450 | U5 | MA1-74183 | ThermoFisher-Scientific | AB6 | 05-212 | Millipore | Plasmid overexpression (Lautz et al 2018) | Smith et al, 2016 |
| 11 | PIKE | AGAP2_MOUSE | Q3UHD9 | 263A | A304-263A | Bethyl | DN8 | 200-401-DN8 | Rockland | X63 Myeloma (Smith et al, 2016) | Smith et al, 2016 |
| 12 | PTEN | PTEN_MOUSE | O08586 | 217702 | MAB847 | Novus Biologicals | 6H2.1 | 04-035 | Millipore | Plasmid overexpression | |
| 13 | RAPTOR | RPTOR_MOUSE | Q8K4Q0 | Polyclonal | 42-4000 | ThermoFisher-Scientific | 514208 | MAB5957 | Novus Biologicals | siRNA | RNAi |
| 14 | Rheb | RHEB_MOUSE | Q921J2 | 344912 | MAB3426 | R&D Systems | JG37-12 | NBP2-75646 | Novus Biologicals | Plasmid overexpression | RNAi |
| 15 | RICTOR | RICTR_mouse | Q6QI06 | 7B3 | 05-1471 | Novus Biologicals | 9F1.2 | Novus NBP1-51645 | Millipore | mouse KO ESC lysate | |
| 16 | TSC1 | TSC1_MOUSE | Q9EP53 | 5C8A12 | 37-0400 | ThermoFisher-Scientific | Polyclonal | NB100-2315 | Novus Biologicals | Mouse Embryonic fibroblast lysate (TSC1(-/-); ) | RNAi |
| 17 | TSC2 | TSC2_MOUSE | Q61037 | 614204 | MAB40401 | R&D Systems | SC05-59 | NBP2-67552 | Novus Biologicals | Plasmid overexpression | RNAi |
